# Uncovering footprints of natural selection through time-frequency analysis of genomic summary statistics

**DOI:** 10.1101/2022.10.05.510997

**Authors:** Sandipan Paul Arnab, Md Ruhul Amin, Michael DeGiorgio

**Affiliations:** Department of Electrical Engineering and Computer Science, Florida Atlantic University, Boca Raton, FL 33431, USA

**Author notes:** Corresponding authors* (S.P.A.), (M.D.).

## Abstract

Natural selection leaves a spatial pattern along the genome, with a distortion in the haplotype distribution near the selected locus that becomes less prominent with increasing distance from the locus. Evaluating the spatial signal of a population-genetic summary statistic across the genome allows for patterns of natural selection to be distinguished from neutrality. Different summary statistics highlight diverse components of genetic variation and, therefore, considering the genomic spatial distribution of multiple summary statistics is expected to aid in uncovering subtle signatures of selection. In recent years, numerous methods have been devised that jointly consider genomic spatial distributions across summary statistics, utilizing both classical machine learning and contemporary deep learning architectures. However, better predictions may be attainable by improving the way in which features used as input to machine learning algorithms are extracted from these summary statistics. To achieve this goal, we apply three time-frequency analysis approaches (wavelet transform, multitaper spectral analysis, and S-transform) to summary statistic arrays. Each analysis method converts a one-dimensional summary statistic arrays to a two-dimensional image of spectral density or visual representation of time-frequency analysis, permitting the simultaneous assessment of temporal and spectral information. We use these images as input to convolutional neural networks and consider combining models across different time-frequency representation approaches through the ensemble stacking technique. Application of our modeling framework to data simulated from neutral and selective sweep scenarios reveals that it achieves almost perfect accuracy and power across a diverse set of evolutionary settings, including population size changes and test sets for which sweep strength, softness, and timing parameters were drawn from a wide range. Moreover, a scan of whole-genome sequencing of central European humans recapitulated previous well-established sweep candidates, as well as predicts novel cancer associated genes as sweeps with high support. Given that this modeling framework is also robust to missing data, we believe that it will represent a welcome addition to the population-genomic toolkit for learning about adaptive processes from genomic data.

## Introduction

A number of phenomena shape genomic diversity, including nonadaptive processes, such as mutation, recombination, genetic drift, and migration as well as adaptive processes, such as positive, negative, and balancing selection [Gillespie, 2004]. Many of these events leave local footprints of altered haplotypic variation across individuals in populations, restructuring the landscape of diversity across the genome [Fay et al., 2001, Prezeworski et al., 2005, Schlamp et al., 2016, Charlesworth, 2006]. To learn about such processes, myriad summary statistics have been developed over decades, providing tools for testing whether patterns in genetic variation match expectations, either from theoretical models or from mean patterns observed from simulations [*e.g*., Tajima, 1983, Garud et al., 2015]. One of the most extensively-studied population-genetic phenomena that has received substantial attention in terms of method development over the past few decades is natural selection.

Natural selection is a process that acts on traits of individuals within an environment, leading to differential fitness among individuals that may result in changes in the frequencies of alleles that code for such traits within a population [Gillespie, 2004]. Genomic studies of a wide range of populations and species have been analyzed using a variety of summary statistic methodologies to search for signatures of natural selection [*e.g*., Glinka et al., 2003, Lucas et al., 2019, Xue et al., 2021]. Summary statistics developed throughout the past several years rely heavily on the haplotype frequency spectrum [*e.g*., Garud et al., 2015], whereas more classical summaries focused more on the site frequency spectrum [*e.g*., Tajima, 1983]. These varied approaches interrogate different aspects of genomic variation, and lend greater ability to detect specific forms of adaptation [Vitti et al., 2013].

However, such summary statistics typically make simplifying assumptions about expected patterns of variation, and can be both underpowered and non-robust to confounding factors when applied individually. To overcome the pitfalls associated with using a single summary statistic to uncover signals of evolutionary processes, combining the knowledge garnered from a plethora of summary statistics has become an emerging trend [Schrider and Kern, 2018]. Specifically, the recent expansion of modeling frameworks that combine sets of measured values to discriminate among diverse evolutionary scenarios is owed to the advancement of computational technologies and resurgence of statistical machine learning and artificial intelligence. These predictive models have been shown to typically offer greater detection power and accuracy, while also combating the drawbacks of individual hand-engineered summary statistics [*e.g*., Lin et al., 2011, Schrider and Kern, 2016, Sheehan and Song, 2016, Sugden et al., 2018, Kern and Schrider, 2018, Mughal and DeGiorgio, 2019, Mughal et al., 2020]. These machine learning techniques employ diverse modeling paradigms, and have differing performances and robustness to confounding factors depending on how the data are modeled as well as the types of summary statistics that are used as input to the models. However, some models suffer from lack of robustness to technical artifacts expected from empirical studies, and all methods show room for improvement in prediction performance.

To glean more information from input summary statistics, many of these models [*e.g*., Lin et al., 2011, Kern and Schrider, 2016, Sheehan and Song, 2016] construct feature sets so that they capture the expected spatial autocorrelation of variation in a local genomic region. That is, input summary statistics are calculated in a number of contiguous genomic windows, with a hope that the machine learning models will place greater emphasis on certain windows and less emphasis on others. However, explicitly modeling these autocorrelations may have the potential for improving prediction performance. As an example, Mughal et al. [2020] developed a method for learning about positive natural selection by utilizing multiple summary statistics computed in contiguous genomic windows as input, and then modeled the autocorrelation across these windows by estimating the underlying continuous functional form of each summary statistic. Specifically, Mughal et al. [2020] employed a spectral analysis technique termed the discrete wavelet transform, which decomposed the summary statistic vectors in the form of multi-level details of constituent low- and high-frequency regions, enabling additional meaningful information to be extracted from the summary statistics.

Spectral analysis of signals has been extensively applied in various domains, including biomedical sciences [O’Brien et al., 2019], power systems [Khan and Pierre, 2018], and seismography [Puryear et al., 2012], to extract information about the source (or process) responsible for the generation of the examined signals from their oscillatory characteristics. One way to extract information from the signal is to divide the signal into time-localized components and examine each part of the signal independently though time-frequency analysis images. Different time-frequency analysis methods focus on different characteristics of a signal [Xiang and Hu, 2012], and thus, images of the characteristics identified by different time-frequency analysis methods can be used as input to established modeling frameworks that are able to extract meaningful information and make accurate predictions. One mechanism for attempting to learn such features is with convolutional neural networks [CNNs, LeCun et al., 1998].

CNNs offer a framework for extracting features from inputs that can be one-dimensional vectors, two-dimensional matrices (or grayscale images), and three-dimensional tensors (or color images) [LeCun et al., 1998]. Several studies have shown the effectiveness of CNNs for detecting evolutionary events for both one- and two-dimensional signals [Flagel et al., 2019, Kern and Schrider, 2016, Torada et al., 2019, Gower et al., 2021]. Indeed, CNNs have been applied in the context of learning about evolutionary processes from image representations of haplotype variation, and have been demonstrated to often have greater power and accuracy compared to the current state-of-the-art summary statistic-based methods [Flagel et al., 2019, Isildak et al., 2021]. A hybrid application of using two-dimensional time-frequency analysis images generated through signal decomposition to train CNNs has the potential to empower the CNNs to make more effective predictive models. To employ this modeling strategy, one-dimensional summary statistic signals need to be converted into two-dimensional time-frequency analysis images [Cohen, 1995, Sejdi et al., 2009], which provide information about the spectral estimates of the underlying source (or process) that generates genomic variation.

Therefore, we seek to improve evolutionary process classifiers, by adding a layer of spectral inference of the underlying process generating the genetic variation. To that end, we use the detection of positive natural selection as a test case, as this setting is where the majority of population-genetic machine learning development has focused, and thus represents a test case for illustrating the performance gains by modeling input data differently. Positive natural selection increases the frequencies of alleles in a population that code for beneficial traits, potentially leading to fixation within the population and ultimately reducing diversity at the selected locus [Gillespie, 2004]. As this beneficial allele increases in frequency, alleles on the same haplotype at nearby neutral loci also increase in frequency through a process known as genetic hitchhiking [Smith and Haigh, 1974]. The resulting loss of haplotypic diversity around the selected locus is known as a selective sweep [Przeworski, 2002, Hermisson and Pennings, 2005], and is a footprint that is often used to uncover signals of past positive selection. Depending on the number of distinct haplotypes that have risen to high frequency, selective sweeps can be categorized as either soft or hard, with hard sweeps typically easier to detect due to their more conspicuous genomic pattern [Przeworski, 2002, Hermisson and Pennings, 2005, Garud et al., 2015].

In this article, we examine the utility of applying three signal decomposition methods on arrays of summary statistics computed over contiguous windows to generate time-frequency analysis images [Daubechies, 1992, Stockwell et al., 1996, Thomson, 1982], and develop machine learning methods trained with these images. We additionally employ ensemble-based stacking procedures [Hastie et al., 2009] that aggregate the results of individual classifiers with the goal of further improving power and accuracy to detect sweeps from genome variation. With this in mind, we introduce an approach termed *SISSSCO* (Spectral Inference of Summary Statistic Signals using COnvolutional neural networks) with open-source implementation available at https://www.github.com/sandipanpaul06/SISSSCO. As an empirical test case, we then apply our trained *SISSSCO* models to whole-genome data of the well-studied central European human individuals sequenced by the 1000 Genomes Project [The 1000 Genomes Project Consortium, 2015]. *SISSSCO* identifies multiple genes, including *LCT, ABCA12, SLC45A2, HLA-DRB6*, and *HCG9*, which have been identified as sweep candidates from previous studies. *SISSSCO* also identified several novel sweep candidates, including *PDPN, WASF2, LRIG2*, and *SDAD1*.

## Results

In this section, we begin by highlighting power and accuracy to detect selective sweeps using various strategies that combine different time-frequency decompositions of summary statistic signals as well as stacking of trained CNN architectures. We also compare the performance of these approaches with other contemporary machine learning methods that take summary statistics as input to detect sweeps. We then investigate how confounding factors, like changing population sizes over time and the existence of missing data, influence predictive accuracy, power, and robustness. Finally, as a proof of concept, we test our new approaches using a genomic dataset from a human population that has been extensively studied.

### Modeling description

To train and test our models, we simulated neutral and sweep replicate observations using the coalescent simulator `discoal` [Kern and Schrider, 2016] under either an equilibrium constant-size demographic history of 10,000 diploid individuals [Takahata, 1993] or under a nonequilibrium history inferred from central European human genomes [Terhorst et al., 2017] that includes a recent severe population bottleneck. Per-site per-generation mutation (*μ* = 1.25 × 10^−8^) and recombination rates (exponential distribution with mean *r* = 10^−8^ and truncated at 3r) were chosen to reflect expectations from human genomes and previous studies [Scally and Durbin, 2012, Payseur and Nachman, 2000, Schrider and Kern, 2016]. For each simulated replicate, we sampled 198 haplotypes of length 1.1 megabase (Mb) to match the number of sampled haplotypes in our empirical experiments.

At the center of simulated sequences for sweep observations, we introduced a beneficial mutation that became selected for at a frequency of *f* ∈ [0.001, 0.1] (drawn uniformly at random on a logarithmic scale) with per-generation selection coefficient *s* ∈ [0.005, 0.5] (drawn uniformly at random on a logarithmic scale) and became fixed in the population *t* generations prior to sampling. For each of the two demographic scenarios, we generated two datasets: one with the sweep completing at time of sampling (*t* = 0) and a setting that should be more difficult to distinguish from neutrality, with *t* ∈ [0,1200] drawn uniformly at random, permitting the processes of mutation, recombination, and genetic drift to erode genomic footprints of the selective sweep after fixation. We denote these four datasets as Equilibrium_fixed, Equilibrium_variable, Nonequilibrium_fixed, and `Nonequilibrium_variable`, where the demographic history is given by either equilibrium (constant-size) or nonequilibrium (European human bottleneck) and the time of sampling after sweep completion is given by either fixed (*t* = 0) or variable (*t* ∈ [0,1200]).

For each class (neutral or sweep), we generated 11,000 independent simulated replicate observations, with 9,000, 1,000, and 1,000 observations reserved for training, validation, and testing. For each replicate, we computed summary statistics across the simulated sequence to obtain nine one-dimensional signals to use as features for downstream modeling identical to the ones used in Mughal et al. [2020] (see *Methods* for summary statistic computation on simulated data). The initial summary statistic that we explored in our model training is the mean pairwise sequence difference (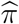; Tajima [1983]) estimated across sampled haplotypes. The dataset containing instances of 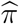 computed as a one-dimensional signal across a genomic sequence of neutral and selective sweep regions was used to test the efficacy of each of the three time-frequency analysis methods, which we discuss in detail within the *Methods* section and briefly summarize in this section.

The two-dimensional images that we obtain by performing time-frequency analysis on a onedimensional signal (*e.g*., 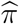) are then fed into a CNN [LeCun et al., 1998], which is depicted in Figure 1. The CNN has an input size of (*N, m, n, c*) containing *N* training observations of *c* different summary statistic signals decomposed as *m* × *n* images through time-frequency analysis. Here we have *N* = 18, 000, *m* = 65, and *n* = 128. As we are currently only considering a single signal based on the 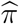 statistic, we are using a *c* =1 channel input for our CNN. The CNN has two convolution layers with 32 filters, kernels of size 3 × 3 [Agrawal and Mittal, 2020], and a stride of two [Kong and Lucey, 2017] with zero padding [Hashemi, 2019]. Each convolution layer is then followed by an activation layer using a rectified linear unit (ReLU), as well as a batch normalizing layer [Goodfellow et al., 2016]. The convolution layers are followed by a dense layer containing 128 nodes, which is the same as the input signal length *n*. The dense layer also contains an elastic-net style regularization penalty [Zou and Hastie, 2005], whereby network weights shrink in magnitude together toward zero through an *L*_2_-norm penalty while simultaneously performing feature selection by setting some weights to zero through an *L*_1_-norm penalty [Hastie et al., 2009]. The fraction of regularization deriving from the *L*_2_-norm penalty is controlled by hyperparameter *α* ∈ {0.0, 0.1, …, 1.0} and the amount of total regularization is controlled by hyperparameter λ ∈ {10^−6^, 10^−5^, …, 10^5^}. The model also utilizes a dropout layer with dropout rate hyperparameter *x* ∈ {0.1, 0.2, …, 0.5} to further prevent model overfitting by reaching a saturation point [Srivastava et al., 2014, Goodfellow et al., 2016]. The model is trained with each (*α, λ, x*) hyperparameter triple, and the best model is chosen as the one with the smallest validation loss, where we employ the categorical cross-entropy loss measurement. We deployed the `keras` Python library [Chollet et al., 2015] for training of CNNs and making downstream predictions from the learned models.

**Figure 1:**
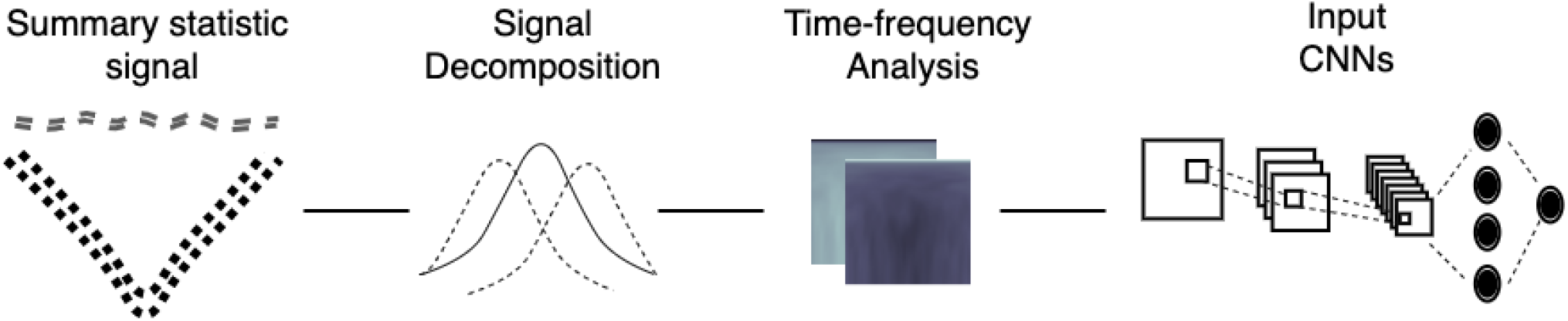
Depiction of a *c* = 1 channel convolutional neural network (CNN) architecture. A summary statistic signal of length *n* = 128 is used as input to a time-frequency analysis method (either wavelet decomposition, multitaper analysis, or S-transform) to decompose the signal into a matrix of dimensions *m × n*, with *m* = 65, which is then standardized at each element based on the mean and standard deviation across all *N* = 18, 000 training observations, and is then used as input to a CNN. The CNN has two convolution layers (three layers for the S-transform), followed by a dense layer with *n* nodes containing both elastic-net and dropout regularization. The output layer of the CNN is a softmax that computes the probability of a sweep.

The first of three time-frequency analysis methods that we consider is wavelet decomposition. Specifically, we assume that each 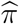 sequence of length *n* =128 represents a sample from a continuous wavelet containing *n* data points. This signal is then decomposed by a level *m* wavelet analysis method, with the Morlet wavelet [Bernardino and Santos-Victor, 2005] selected as the mother wavelet. Level *m* = 65 is chosen for the scalograms generated to match the size of the periodograms that result from the other two time-frequency analysis methods that we subsequently introduce. Every decomposed signal generates an *m* × *n* dimensional scalogram matrix. A more detailed treatment of the wavelet decomposition for time-frequency analysis is provided in the *Methods* section, and we employed the `PyWavelets` Python package [Lee et al., 2019] to construct scalogram images.

Next, for the multitaper time-frequency analysis approach, to derive the periodogram of the estimate of the true power spectral density from a wavelet array of size *n* =128 using the multitaper spectral analysis method, we used a window length of *n*. We calculated discrete prolate spheroidal sequence (DPSS) tapers over time-half bandwidth parameter (*n* × Δ*f*/2) values in {2, 2.5, …, 4} and a DPSS window size of *m* = *n*/2 + 1 = 65, which results in a matrix of tapering windows of size *m* × *n* and a vector of eigenvalues of length *m*. Here, Δ*f* is the bandwidth of the most dominant frequencies in the frequency domain such that *n* × Δ*f*/2 > 1Hz. Using this matrix and vector, a periodogram of size *m* × *n* is generated, which is the same as the dimension of the scalogram that we considered with the wavelet analysis method. See the *Methods* section for a complete detailed description of multitaper analysis. We utilized the `spectrum` Python package [Cokelaer and Hasch, 2017] to generate multitaper periodogram images.

Finally, for time-frequency analysis using the Stockwell transform (also known as the S-transform) we used the same datasets as the previous two time-frequency analysis approaches. The S-transform returns a periodogram matrix estimate of the true power spectral density that has size *m* × *n*, where *m* = *n*/2 + 1 and where the length of the signal is *n* = 128. The periodogram has the same image size as the previous two methods. See the *Methods* section for further details on the S-transform. We used the `stockwell` Python package [Satriano, 2017] to estimate S-transform periodogram images. The images are then fed into a CNN with identical architecture to that of the previous two methods with the addition of a third convolution layer, which we included as we found that adding this extra convolution layer substantially increased performance under the S-transform image inputs.

### Effect of signal decomposition

Figure S1 presents heatmaps of the raw time-frequency images, averaged across simulated replicates, for neutral and selective sweep regions using three signal decomposition methods. However, based on these raw images, it is difficult to visually distinguish between sweeps and neutrality for each of the time-frequency analysis methods. To better explore the visual differences within these matrices, we scaled each element of each time-frequency analysis matrix to have unit standard deviation across the neutral and sweep replicates. The mean scaled matrices depicted in Figure 2 show the emergence of more-readily distinguishable patterns between sweeps and neutrality. The wavelet decomposition results display a clear distinction between the two classes, with a triangular bulge in the mid-segment of the sweep scalogram that is not present within the neutral scalogram. This pattern indicates that the selective sweep signals have information in the middle windows between windows 45 and 85 that is not present in neutral signals. The mean periodograms generated by multitaper analysis depict a rib-cage like structure in the mean sweep periodogram, whereas the mean sweep periodogram generated by S-transform shows a T-shaped construct in the mid-portion of the image.

**Figure 2:**
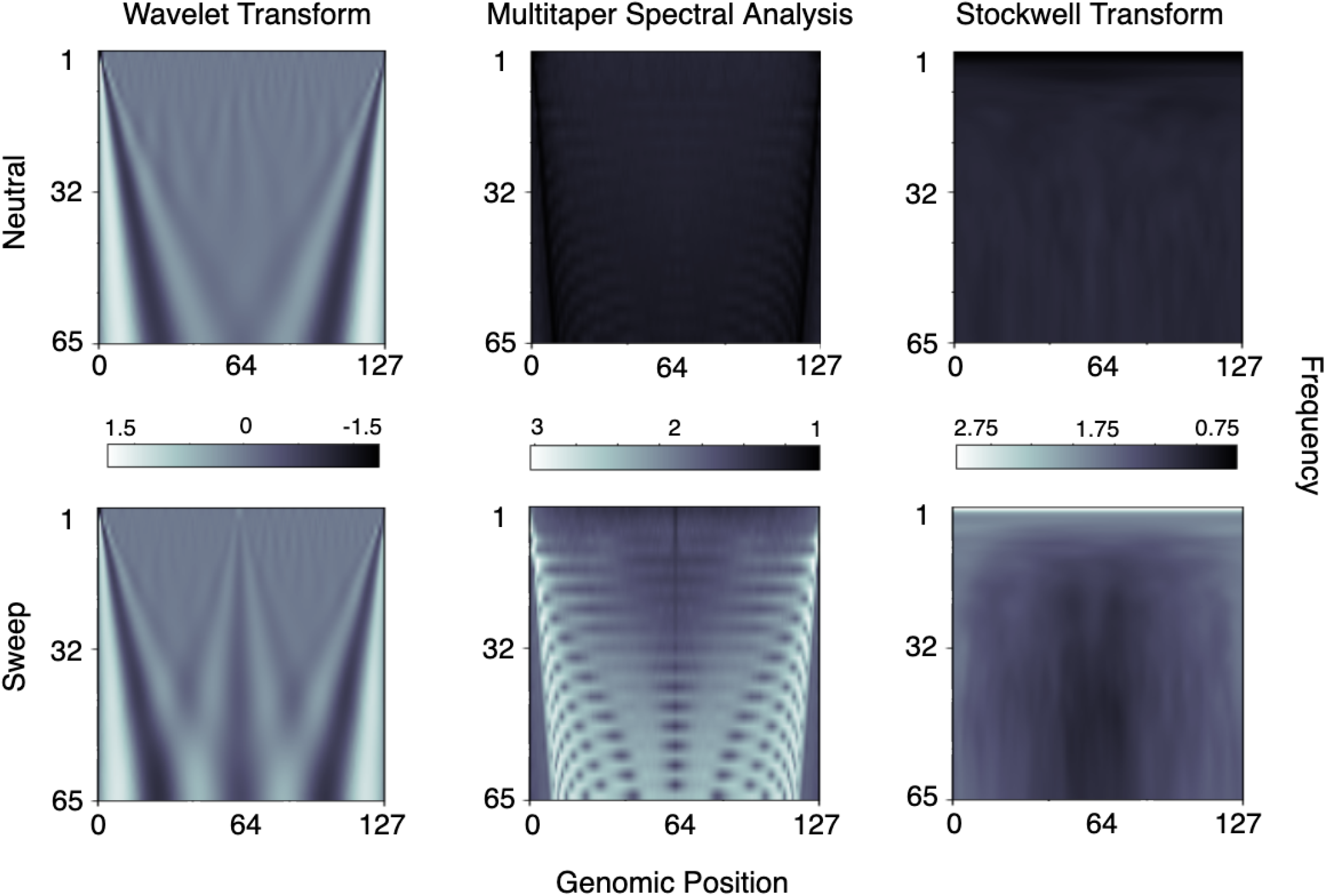
Mean time-frequency analysis input matrices for *n* = 128 windows of the mean pairwise sequence differences 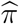 across the *N*/2 = 9, 000 neutral and *N*/2 = 9, 000 sweep replicates under the `Equilibrium_fixed` dataset containing an equilibrium constant-size demographic history and a sweep that completed *t* = 0 generations before sampling. Top row are neutral simulations and bottom row are sweep simulations. Time-frequency methods are depicted from left to right columns for the wavelet decomposition, multitaper analysis, and the S-transform, respectively. Elements of each matrix have been scaled to have a standard deviation of one across all *N* simulated replicates for a given time-frequency analysis method.

**Figure 3:**
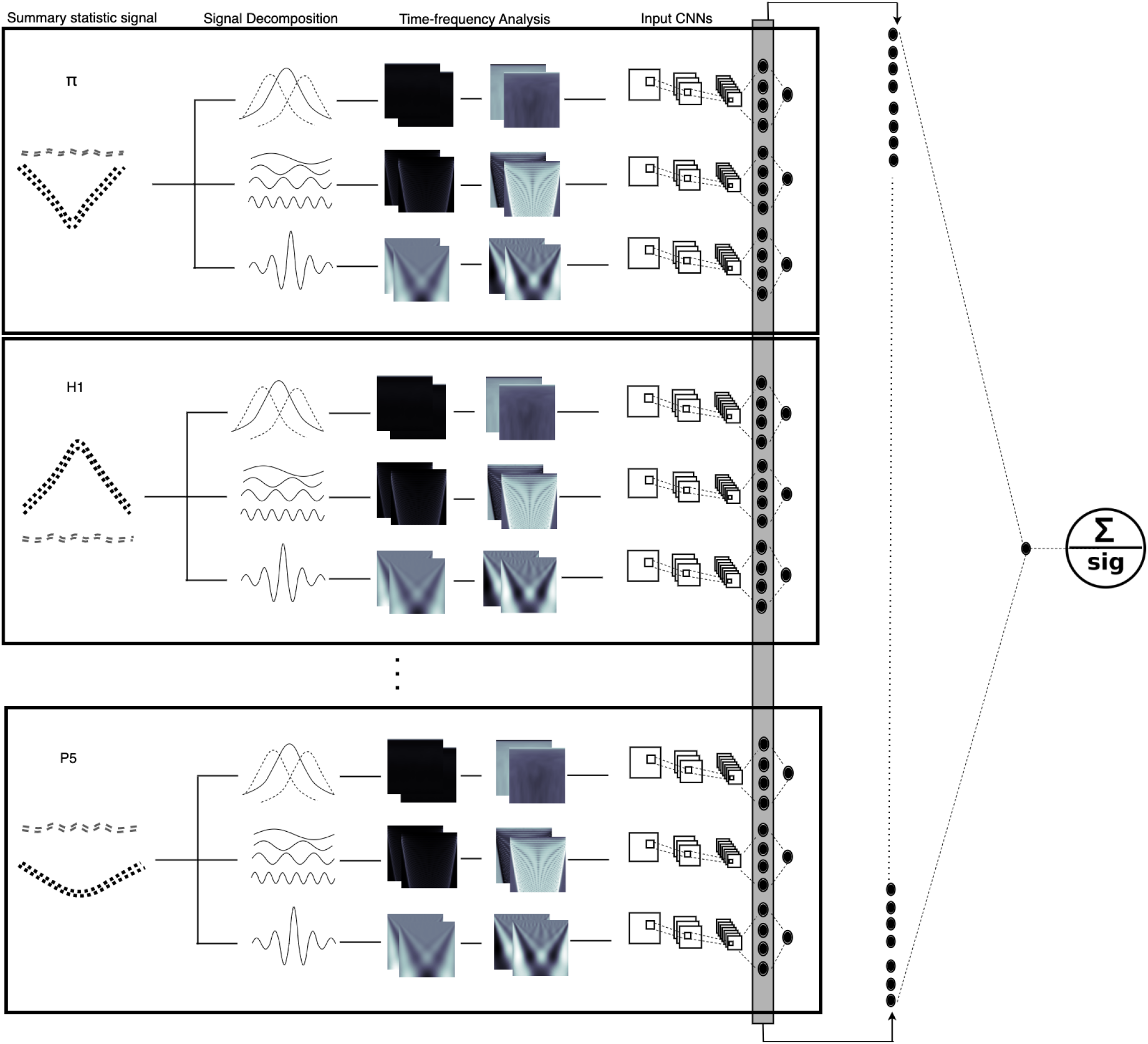
Depiction of the *SISSSCO*[*27CD*] model. Each summary statistic signal (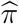, *H*_1_, *H*_12_, *H*_2_/*H*_1_ and frequencies of the first five most common haplotypes respectively denoted by *P*_1_ to *P*_5_) of length *n* = 128 is used as input to each of the three time-frequency analysis method (wavelet decomposition, multitaper analysis, and S-transform) to decompose the signal into three matrices of dimension *m* × *n*, with *m* = 65, which are then each standardized at each element based on the mean and standard deviation across all *N* = 18, 000 training observations. These 27 images (nine statistics across three time-frequency analysis methods) each used as input to train 27 independent convolutional neural networks (CNNs). The CNNs have two convolution layers (three layers for the S-transform), followed by a dense layer with *n* nodes containing both elastic-net and dropout regularization. The output layer of the CNN is a softmax that computes the probability of a sweep. After training, the model parameters are fixed, and the dense layers of the 27 CNNs are concatenated and these 27*n* = 3, 456 nodes are used as input to a new output layer, which computes the probability of a sweep as a softmax.

The standardized (combined centering and scaling) images in Figure S2 that are ultimately used as input to CNNs show that the classes can be easily visually differentiated as the images show exactly opposite patterns for the two classes, with the images for neutral regions having lower values for the majority of the area in the images. These opposite patterns are due to centering. A peach pit shape is present in the center of both mean sweep and neutral periodograms generated by the S-transform, albeit represented by two distinctly different shades corresponding to positive and negative values, respectively. The rib-cage structure is also present in mean periodograms of both classes in the images created by multitaper analysis, with different shades for the two classes corresponding mostly to positive and negative values.

Figures 2, S1, and S2 highlight the qualitative patterns in images derived from neutral and sweep settings that result from three different time-frequency analysis methods applied to a sequence of 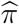 values calculated across contiguous genomic windows. Given that these images show qualitative differences between sweeps and neutrality, our goal is to evaluate the predictive ability of discriminating between sweeps and neutrality from such input images. Therefore, using the CNN architecture described above in the *Modeling description* subsection, we fed the images derived upon application of the time-frequency analysis methods to a sequence of 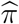 values to evaluate classification rates and accuracy. Figure S3 shows that the models trained on wavelet analysis scalogram and S-transform periodogram images have an imbalance in their classification rates, with skews toward detecting neutral regions more accurately than the sweep regions. In contrast, the model trained on multitaper analysis periodogram images with a time half-bandwidth parameter of 2.5 displays greater accuracy for correctly estimating sweeps compared to neutral regions, whereas changing the time half-bandwidth parameter to 2.0 or lower results in classification rates more skewed toward correctly detecting neutrality. Because we want to avoid false discoveries of sweeps, higher time half-bandwidth parameter values are more expensive computationally, and time half-bandwidth parameters higher than 2.5 did not change performance significantly in our preliminary tests, we selected 2.0 for future multitaper experiments.

### Stacking models to enhance sweep detection

We have three models trained with three signal decomposition methods that have yielded comparable but slightly differing results (Figure S3). We now discuss architectures to increase the learning capacity of our models when trained to jointly consider all three time-frequency analysis images. Our previous experiments explored a single summary statistic signal 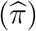 to decompose and train the models with time-frequency analysis images. Following Mughal et al. [2020], we next compute nine one-dimensional summary statistic signals (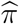, *H*_1_, *H*_12_, *H*_2_/*H*_1_ and frequencies of the five most common haplotypes) per simulated replicate and generate nine time-frequency analysis images for each of the three time-frequency analysis methods, resulting in 27 different images.

The first joint modeling approach taken was to train three separate models using three signal decomposition methods with nine images per replicate provided as input to a CNN, with one image for each of the *c* =9 channels of the CNN (Figure S4). These models were then concatenated and trained in three different ways. The first of these three strategies is to train each of the three nine-channel CNNs, fix the weights of the trained CNNs, and concatenate their output layers (sweep probability values) into a three-element vector of sweep probabilities. The linear combination of these sweep probabilities is then used as input to a new softmax function to predict the probability of a sweep from evidence of the three pretrained CNNs. The final weights of the linear combination leading to the new softmax function are trained, and we denote this method by *SISSSCO*[*3CO*] (three input CNNs and concatenation of the output layer). The weights of the three individually trained CNNs are not retrained in the final model. A depiction of this *SISSSCO*[*3CO*] architecture is given in Figure S5. In the next strategy, we instead concatenated the dense layers of the three nine-channel CNNs, leading to a vector of 3 × 128 = 384 elements that we send to a new softmax layer as in the *SISSSCO*[*3CO*] method. As with *SISSSCO*[*3CO*], we trained the weights of the linear combination leading from the concatenation of the dense layers to the new softmax function, but did not retrain the weights of the three individually trained CNNs, and we denote this method by *SISSSCO*[*3CD*] (three input CNNs and concatenation of the dense layer). A depiction of the *SISSSCO*[*3CD*] architecture is given in Figure S6. The third and final strategy, has an identical architecture of the *SISSSCO*[*3CD*] model, with one key difference—the weights of the entire concatenated model are jointly trained. We denote this method by *SISSSCO*[*3MD*] (three input CNNs and merging of the dense layer prior to training). A depiction of the *SISSSCO*[*3MD*] architecture is given in Figure S7.

The second joint modeling approach is more complex than the first. Specifically, we construct three CNNs per summary statistic based on the three signal decomposition methods, resulting in 27 distinct CNNs each with *c* =1 channel (Figure 1). Similar to the previous concatenation strategies, the concatenation and training were accomplished in an identical fashion by pretraining individual CNNs and concatenating output layers (model denoted by *SISSSCO*[*27CO*]), pretraining individual CNNs and concatenating dense layers (model denoted by *SISSSCO*[*27CD*]), and concatenating dense layers of individual CNNs with all weights in the subsequent merged model trained (model denoted by *SISSSCO*[*27MD*]). Both *SISSSCO*[*27CD*] and *SISSSCO*[*27MD*] methods result in the most complex final models, with the dense layer containing 128 × 27 = 3, 456 nodes. Though *SISSSCO*[*27CD*] and *SISSSCO*[*27MD*] have the same number of concatenated dense layer nodes, the node weights are not set prior to concatenation for *SISSSCO*[*27MD*], making *SISSSCO*[*27MD*] the most computationally expensive method among all the six models. The architectures of the *SISSSCO*[*27CO*], *SISSSCO*[*27CD*], and *SISSSCO*[*27MD*] models are depicted in Figures S8, 3, and S9, respectively. In the next subsection, we evaluate the accuracies and powers of the six *SISSSCO* models on idealistic constant-size demographic history datasets.

### Power and accuracy to detect sweeps

All of our six *SISSSCO* models have high classification accuracies and powers on the two constant-size demographic history datasets (Figures S10-S13). Of these, *SISSSCO*[*27CD*] exhibited uniformly highest accuracy to discriminate sweeps from neutrality, reaching 99.75 and 99.80% accuracy on the Equilibrium_fixed and `Equilibrium_variable` datasets, respectively (Figures S10 and S12). However, even the worst performing *SISSSCO* model had high accuracy on each dataset, with *SISSSCO*[*3CD*] achieving an accuracy of 96.50 and 95.45% on the Equilibrium_fixed and `Equilibrium_variable` datasets, respectively (Figures S10 and S12). This lower classification accuracy of *SISSSCO*[*3CD*] compared to the other *SISSSCO* models appears to be primarily driven by a skew in misclassifying neutral regions as sweeps (Figures S10 and S12).

The accuracy results are also reflected in the high powers of the *SISSSCO* models to detect sweeps based on receiver operating characteristic (ROC) curves (Figures S11 and S13). Specifically, *SISSSCO*[*27CD*] achieves an area under the ROC curve of close to one for both datasets (Figures S11 and S13), suggesting that it has perfect power to detect sweeps for even small false positive rates. Moreover, consistent with *SISSSCO*[*3CD*] having the lowest accuracy among the six *SISSSCO* models, the ROC curves show that *SISSSCO*[*3CD*] reaches high power for low false positive rates, but plateaus at this level until high false positive rates (Figures S11 and S13), reducing the overall area under the ROC curve compared to the other *SISSSCO* models. The results show that, though all *SISSSCO* models have high powers and accuracies for sweep detection, the most parameter rich (yet not most computationally expensive) *SISSSCO*[*27CD*] model outperforms all others developed here on the constant-size demographic history datasets (Figures S10-S13).

### Performance relative to comparable methods

We tested the classification performance of our models against three state-of-the-art methods that employ summary statistics as input: *SURFDAWave* [Mughal et al., 2020], diploS/HIC [Schrider and Kern, 2016], and evolBoosting [Lin et al., 2011]. *SURFDAWave* is a wavelet-based classification method that takes as input nine summary statistic arrays, exactly the ones that we have used for our study, and learns the functional form of the spatial distribution of each summary statistic using a wavelet basis expansion to represent the autocorrelation among a summary statistic across the genome. The method then uses estimated wavelet coefficients as input to elastic net logistic regression models for classifying selective sweeps and predicting adaptive parameters.

On the other hand, to detect selective sweeps, diploS/HIC takes a complementary deep learning approach to extract additional information from arrays of different features of population genetic variation. In particular, the deep CNN classifier used in diploS/HIC takes images of a set of multidimensional summary statistic vectors calculated in 11 windows, with the central window denoted as the target. The set of summary statistics considered is different from *SURFDWave*, instead employing a set of summary statistics that assesses nucleotide and multilocus genotype variation without the need for phased haplotypes.

Furthermore, evolBoosting also uses arrays of different summary statistics as input and applies boosting to detect selective sweeps from neutrality. The purpose of the boosting [Schapire, 1999] ensemble technique is to create an optimum combination of simple classification rules obtained from the base classifiers [Hastie et al., 2009], which are themselves quite simple and not particularly accurate. This strategy is inspired by the observation that, in most cases, an ensemble of basic rules can outperform classifiers individually [Schapire, 1999]. Boosting involves fitting data instances to a model, and training the model in a series. Incorrect predictions are used to train a subsequent model. Each newly added base model improves prediction error by accounting for error that was not captured by the set of prior base models. At each iteration, the less reliable rules of each base classifier are aggregated into a single, more reliable rule.

These three methods consider both linear and nonlinear classification strategies, with *SURFDAWave* employing a linear model and diploS/HIC and evolBoosting nonlinear approaches. We applied these three methods using their default settings, such as window lengths, window sizes, sets of features, and summary statistic generation and usage. It is important to note that diploS/HIC was originally developed to discriminate among five classes: soft sweeps, hard sweeps, linked soft sweeps, linked hard sweeps, and neutrality. As in Mughal et al. [2020], we retooled the method as a binary classifier to distinguish selective sweeps from neutrality given input summary statistics.

On both the `Equilibrium_fixed` and `Equilibrium_variable` datasets, *SURFDAWave*, diploS/HIC, and evolBoosting achieved relatively high accuracy to discriminate sweeps from neutrality, with the lowest of them (evolBoosting) achieving an accuracy of 97 and 95% on the `Equilibrium_fixed` and `Equilibrium_variable` datasets, respectively (Figures S10 and S12). Similarly, *SURFDAWave* had highest accuracy among the three methods on each dataset, achieving an accuracy of 97.95 and 97.60% on the `Equilibriu_fixed` and `Equilibrium_variable` datasets, respectively (Figures S10 and S12). The marginally lower accuracies of evolBoosting and diploS/HIC compared to *SURFDAWave* appears to be due to an imbalance in their predictions, with extremely high accuracy at correctly classifying neutrality coupled with elevated misclassification rates of sweeps as neutral (Figures S10 and S12). However, this skew toward misclassifying sweeps as neutral is conservative, and is substantially more desirable than a skew toward falsely discovering neutral regions as sweeps. Moreover, as expected, each method had a decrease in accuracy on the more challenging `Equilibrium_variable` dataset (Figure S12) relative to the `Equilibriu_fixed` dataset (Figure S10). In comparison with *SISSSCO*, four of the *SISSSCO* models had higher accuracy than the competing methods on the `Equilibriu_fixed` dataset (Figure S10), whereas three of them showed higher accuracy on the `Equilibrium_variable` dataset (Figure S12).

In terms of method power, *SURFDAWave*, evolBoosting, and diploS/HIC tended to exhibit marginally lower power than the *SISSSCO* models, yet generally still achieved similarly high levels of the area under the ROC curves as *SISSSCO* models on both datasets (Figures S11 and S13). An exception is evolBoosting, which displayed substantially lower area under the ROC curve compared to other methods, achieving a power (true positive rate) close to one for false positive rates close to 0.2, whereas all other methods attained power close to one for false positive rates less than 0.05. These results suggest that under the constant-size demographic history and selection setting explored here, several *SISSSCO* models had higher classification accuracies and powers compared to other leading machine learning methods that use as input summary statistics for detecting sweeps. Moreover, the *SISSSCO*[*27D*] model achieves near perfect classification accuracy and power.

### Influence of population size changes

Our prior experiments have highlighted the excellent classification accuracies and powers for the *SISSSCO* models. However, such test settings were idealistic, in which there has been no demographic changes over time—in contrast to the expectation for real populations. We therefore trained and tested our models on a demographic history estimated from the well-studied human central European population (CEU) from the 1000 Genomes Project dataset [The 1000 Genomes Project Consortium, 2015], for which there is extensive evidence of severe population size changes in recent history [Terhorst et al., 2017].

As with the idealistic constant-size demographic histories, we trained our methods on the `Nonequilibrium_fixed` and `Nonequilibrium_variable` datasets, which differ by whether the time that the sweep completed was fixed at *t* = 0 generations before sampling or variable and drawn from a distribution *t* ∈ [0,1200] generations in the past, respectively. The latter dataset represents a setting that should be more difficult, as it leads to blurring of the boundaries between the sweep and neutral classes. Moreover, we deployed the six *SISSSCO* models as well as the comparison methods (*SURFDAWAave*, diploS/HIC, and evolBoosting) with identical architectures, training paradigms, and quantity of train, test, and validation data as for the constant population size experiments.

Similarly to the constant-size setting, *SISSSCO*[*27CD*] displayed near perfect accuracy of 99.9 and 99.5% to discriminate sweeps from neutrality on the `Nonequilibrium_fixed` and `Nonequilibrium_variable` datasets, respectively (Figures 4 and S14). *SISSSCO*[*27D*] also had uniformly highest accuracy across all tested *SISSSCO* and non-*SISSSCO* methods (Figures 4 and S14). Of the non-*SISSSCO* methods, highest accuracy was achieved by *SURFDAWave* (98.65%), and lowest by evolBoosting of (94.50%) on the `Nonequilibrium_fixed` dataset (Figures S14). On the `Nonequilibrium_variable` dataset we see the same pattern among the non-*SISSSCO* methods, with *SURFDAWave* achieving the highest accuracy (96.55%), and evolBoosting the lowest (93.00%) (Figure 4).

**Figure 4:**
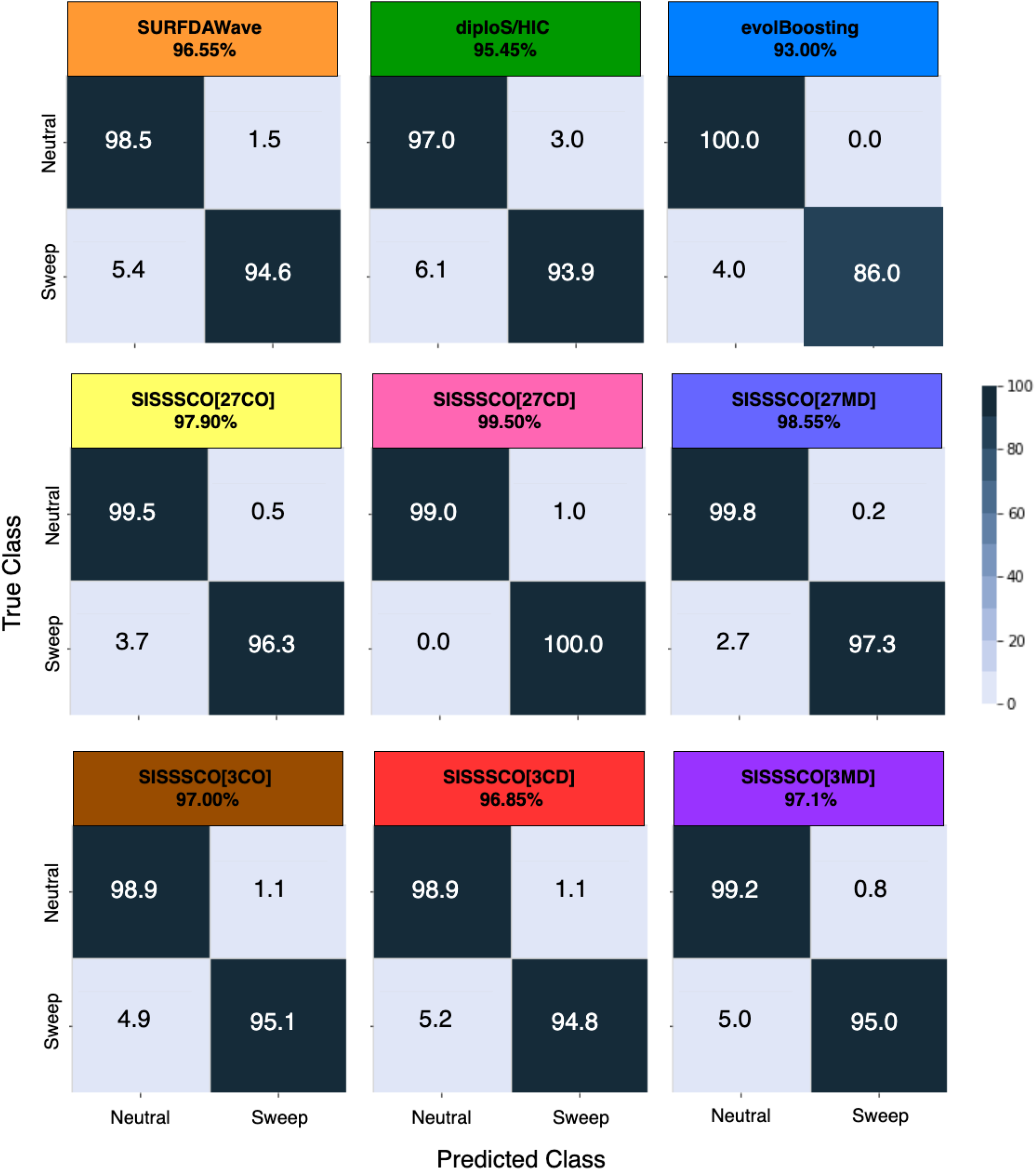
Classification rates and accuracy as depicted by confusion matrices to differentiate sweeps from neutrality on the `Nonequilibrium_variable` dataset for the six *SISSSCO* architectures compared to *SURFDAWave*, diploS/HIC, and evol-Boosting. The `Nonequilibrium_variable` dataset is based on the nonequilibrium recent strong bottleneck demographic history of central European humans (CEU population in the 1000 Genomes Project) and a sweep that completed *t* ∈ [0,1200] generations before sampling.

The high classification accuracies on these datasets are echoed by their high powers to detect sweeps, with all methods aside from evolBoosting achieving areas under the ROC curves that are close to one on the `Nonequilibrium_fixed` dataset (Figure S15). However, the `Nonequilibrium_variable` dataset was more challenging, with *SISSSCO*[*27CD*] the only method achieving near perfect area under the ROC curve, though *SISSSCO*[*27MD*] is close (right panel of Figure 5). For small false positive rates of less than 0.05, evolBoosting has the lowest power, followed by diploS/HIC and *SURFDAWave* having comparable powers, which have lower powers than the three-input *SISSSCO* models (*SISSSCO*[*3CO*], *SISSSCO*[*3CD*], and *SISSSCO*[*3MD*]), with the 27-input *SISSSCO* models (*SISSSCO*[*27CO*], *SISSSCO*[*27CD*], and *SISSSCO*[*27MD*]) harboring the highest overall powers (right panel of Figure 5). The decreased powers of some of the methods are reflected in the imbalance in classification rates demonstrated in Figure 4, for which some methods have a skew toward misclassifying sweeps as neutral. However, as discussed for the constant-size demographic history results, such classification is conservative, as we wish to avoid the alternative skew toward false discovery of sweeps. Overall, our experiments point to *SISSSCO*[*27CD*] having near perfect accuracy and power on the two selection regimes simulated under the nonequilibrium recent strong population bottleneck demographic history.

**Figure 5:**
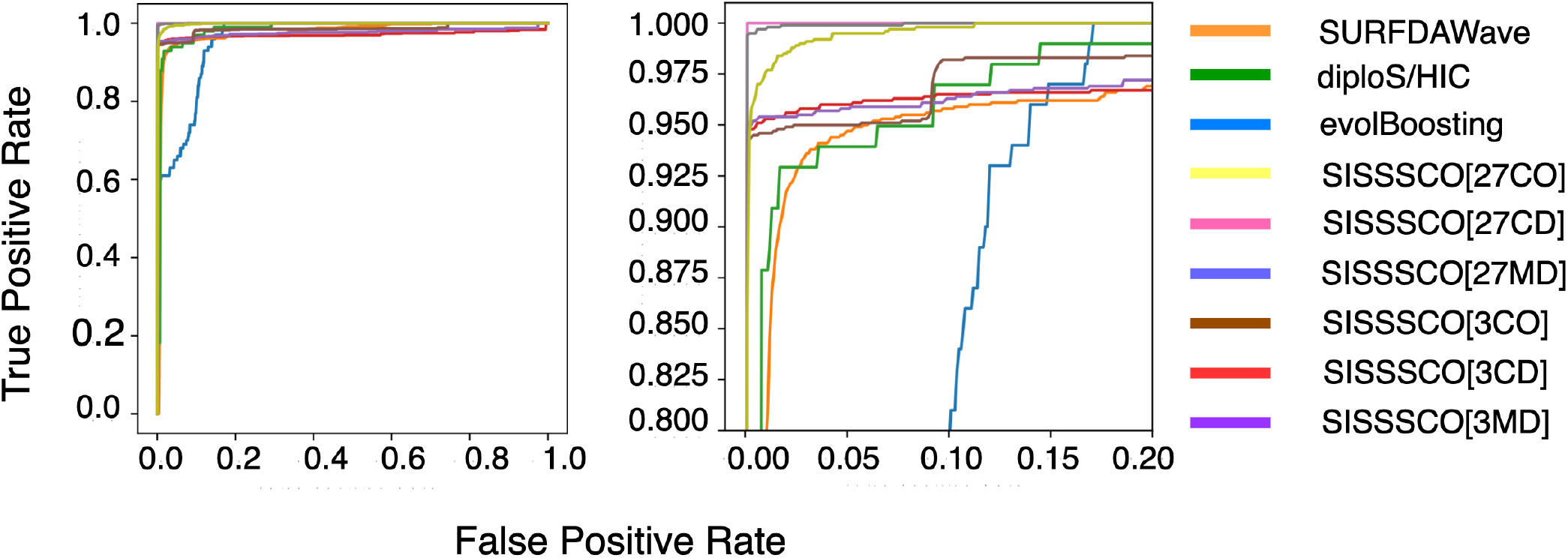
Power to detect sweeps as depicted by receiver operating characteristic curves on the `Nonequilibrium_variable` dataset for the six *SISSSCO* architectures compared to *SURFDAWave*, diploS/HIC, and evolBoosting. The `Nonequilibrium_variable` dataset is based on the nonequilibrium recent strong bottleneck demographic history of central European humans (CEU population in the 1000 Genomes Project) and a sweep that completed *t* ∈ [0,1200] generations before sampling. The right panel is a zoom in on the upper left-hand corners of the left panel.

### Robustness to missing data

The presence of missing genomic segments result from technical artifacts, and can lead to reductions in haplotypic diversity due to unobserved polymorphism. As such losses of local genomic variation can masquerade as selective sweep footprints, missing genomic data may mislead methods that detect sweeps to falsely classify neutral genomic regions harboring missing data as having undergone positive selection. Hence, our goal is to examine whether missing genomic segments within neutrally-evolving test regions lead *SISSSCO* and non-*SISSSCO* methods to falsely identify them as selective sweeps, and whether such missing data hampers the ability of the methods to discriminate between sweeps and neutrality. We therefore simulated an independent set of `discoal` [Kern and Schrider, 2016] replicates for neutral and sweep regions, and generated missing data from these new simulations. Specifically, we followed the protocol of Mughal et al. [2020] by excluding approximately 30% of the SNPs in each simulated replicate, distributed evenly across 10 non-overlapping genomic blocks containing approximately 3% of the SNPs in the replicate. The locations of these blocks are chosen uniformly at random, with a new location chosen for a block if it intersects with locations of previously-placed blocks. We then computed summary statistics using the remaining 70% of SNPs in each replicate, with these statistics measured identically as for the training set using *n* = 128 overlapping windows with a window length of 10 SNPs and a stride of three SNPs calculated over the 400 central SNP sites (200 to the left of the sequence center, and 200 to the right). These one-dimensional summary statistic arrays are then used to generate time-frequency analysis images through the three signal decomposition methods to produce the test dataset consisting of sweep and neutral regions with missing data.

Because the `Nonequilibrium_variable` dataset is the most complex and features a realistic demographic history, we sought to evaluate robustness to missing data on this dataset. We employ models from previous analyses that are trained without missing data (Figures 4 and 5) to these test datasets that contain missing data. As would be expected, the inclusion of missing genomic segments in the test dataset leads to a reduction in classification accuracy across all methods (Figure 6) compared to no missing data (Figure 4). Most notably, diploS/HIC, *SISSSCO*[*3MD*], and evolBoosting experienced moderate to large reductions in accuracy to discriminate sweeps from neutrality, with reductions of 3.85, 4.40, and 5.00%, respectively (compare Figures 4 and 6). This reduction in accuracy appears to be primarily driven by an increase in misclassifying neutral regions as sweeps (Figure 6), for which evolBoosting displays a 23% misclassification rate of falsely detecting neutral regions as sweeps. Of the nine methods compared, *SISSSCO*[*27MD*] has the highest and near perfect accuracy on missing data of 99.95%, exceeding the classification performance of the *SISSSCO*[*27CD*] model that achieved accuracy of 99.50% without missing data but has only 97.90% with missing data. Even on this challenging dataset, *SISSSCO*[*27CD*] and *SISSSCO*[*27MD*] have near perfect powers as evidenced by their near perfect areas under the ROC curves (Figure 7). Therefore, the *SISSSCO*[*27CD*] and *SISSSCO*[*27MD*] models perform comparably well on missing data in terms of power, with *SISSSCO*[*27MD*] edging out *SISSSCO*[*27CD*] in terms of accuracy even though both methods exhibit high accuracy.

**Figure 6:**
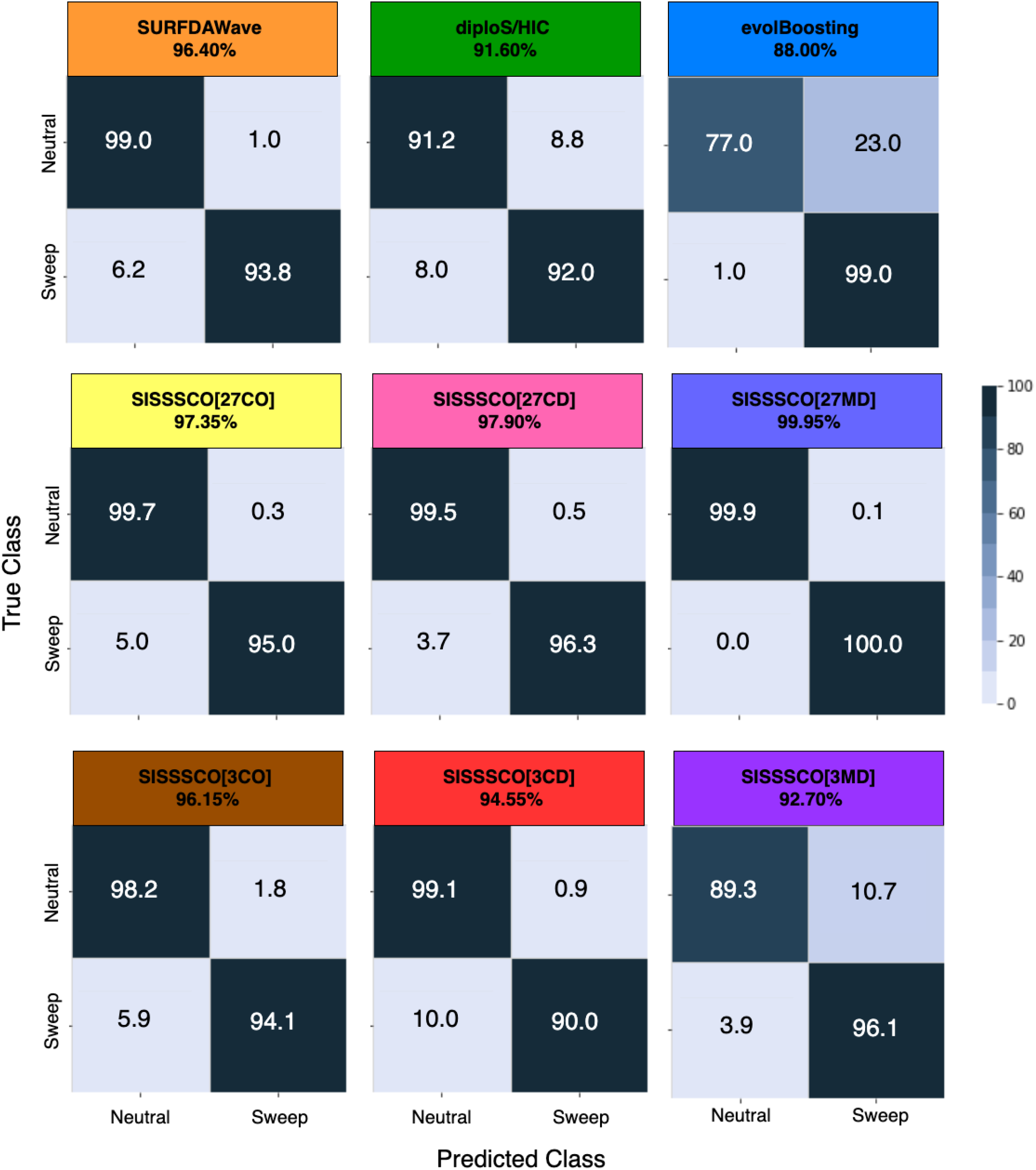
Classification rates and accuracy as depicted by confusion matrices to differentiate sweeps from neutrality on the `Nonequilibrium_variable` dataset when test data contain missing genomic segments for the six *SISSSCO* architectures compared to *SURFDAWave*, diploS/HIC, and evolBoosting. The `Nonequilibrium_variable` dataset is based on the nonequilibrium recent strong bottleneck demographic history of central European humans (CEU population in the 1000 Genomes Project) and a sweep that completed *t* ∈ [0,1200] generations before sampling. Trained models are identical to those in Figure 4 and fitted to training observations without missing data, but the test observations derive from sequences containing approximately 30% missing SNPs distributed evenly across 10 nonoverlapping segments.

**Figure 7:**
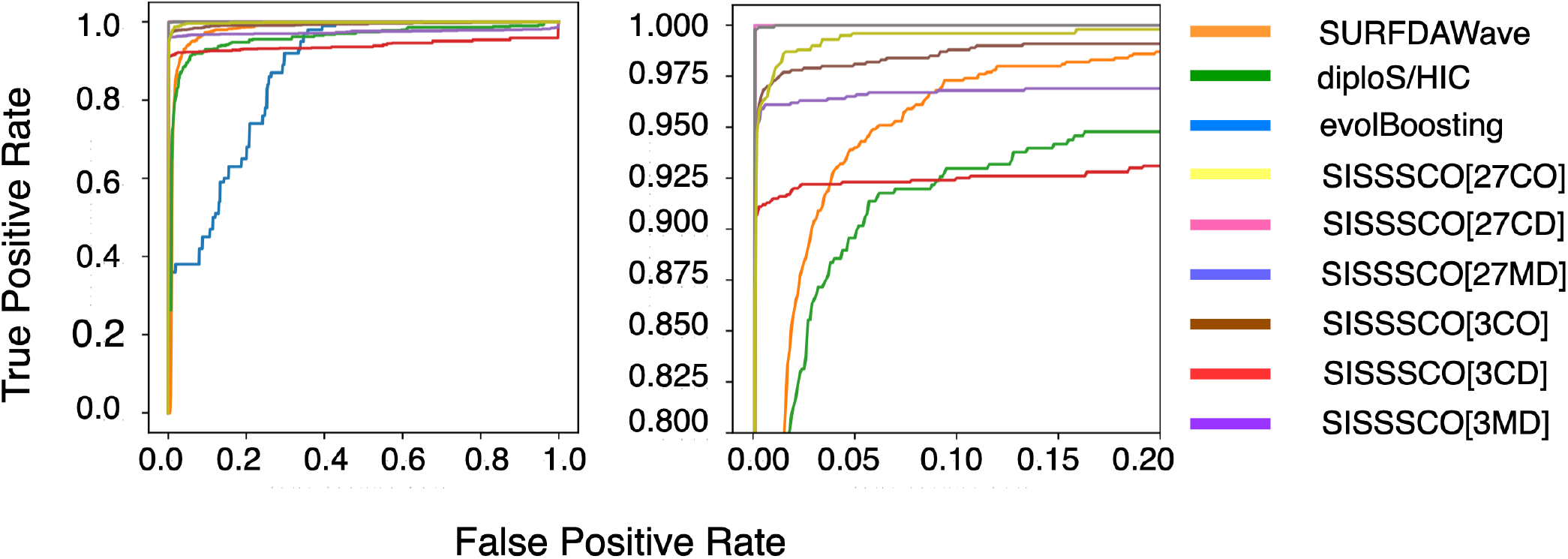
Power to detect sweeps as depicted by receiver operating characteristic curves on the `Nonequilibrium_variable` dataset when test data contain missing genomic segments for the six *SISSSCO* architectures compared to *SURFDAWave*, diploS/HIC, and evolBoosting. The `Nonequilibrium_variable` dataset is based on the nonequilibrium recent strong bottleneck demographic history of central European humans (CEU population in the 1000 Genomes Project) and a sweep that completed *t* ∈ [0,1200] generations before sampling. The right panel is a zoom in on the upper left-hand corners of the left panel. Trained models are identical to those in Figure 5 and fitted to training observations without missing data, but the test observations derive from sequences containing approximately 30% missing SNPs distributed evenly across 10 nonoverlapping segments.

### Application to human genomic data

Until now, we assessed the six *SISSSCO* methods on a number of simulated settings, and compared the results with three competing state-of-the-art methods. Across these tests, *SISSSCO*[*27CD*] was the most consistent performer throughout the evaluation process (Figures 4–7), with a heavier computational cost compared to some of the other *SISSSCO* architectures apart from *SISSSCO*[*27MD*]. Because of its favorable behavior on simulated settings, we apply *SISSSCO*[*27CD*] to variant calls and phased haplotypes of 99 individuals in the CEU population from the 1000 Genomes Project [The 1000 Genomes Project Consortium, 2015] to uncover sweeps in a well-studied human dataset as a proof of concept application of *SISSSCO*.

*SISSSCO* classified most of the genome (approximately 95.5%; Table 1) as neutral, with a mean sweep probability of 17.89%. Increasing the probability threshold for calling sweeps from 0.5 to 0.9 raises the neutral detection rate above 97% (Table 1). We anticipate that the vast majority of the genome will be influenced by non-sweep processes such as background selection and neutral processes, both of which are less likely to produce prominent signatures of low haplotypic variation [Charlesworth et al., 1993, Charlesworth, 2012, Enard et al., 2014]. Based on these properties, we set the sweep footprint detection criterion as a mean prediction probability of at least 0.9 for a set of 10 consecutive prediction windows. Calling sweep regions in this manner circumvents the few isolated data points with marginally high sweep prediction probabilities, and ensures that we are finding peaks in genomes with high sweep support. Figure 8 displays sweep prediction probabilities as a function of genomic position, using a 10-point moving average to generate smoothed curves that match our sweep detection criterion. Of the 22 human autosomes, only the first six contained regions that satisfied our detection criterion, resulting in 20 identified sweep regions containing 22 genes (Table 2 and Figure 8). Among these 22 genes, many are expected from prior scans of European human genomes (*e.g*., *LCT, *ABCA12*, *SLC45A2*, HCG9*, and *HLA-DRB6*), with a few (*e.g*., *PDPN, WASF2, LRIG2, SDAD1, POMGNT1, UQCRH, UPK4*, and *TMPRS11D*) identified as novel candidates in our study.

**Table 1:** Percentage of windows classified as sweep based on sweep probability threshold of 0.5, 0.7, and 0.9 for each of the 22 autosomes of CEU individuals from the 1000 Genomes Project dataset.

| Chromosome | Threshold = 0.5 | Threshold = 0.7 | Threshold = 0.9 |
| --- | --- | --- | --- |
| 1 | 5.31 | 4.31 | 3.32 |
| 2 | 6.59 | 6.41 | 5.89 |
| 3 | 6.60 | 4.13 | 2.55 |
| 4 | 5.56 | 3.89 | 2.92 |
| 5 | 5.99 | 4.76 | 2.12 |
| 6 | 5.19 | 3.72 | 3.01 |
| 7 | 5.71 | 3.92 | 2.02 |
| 8 | 5.00 | 3.67 | 2.01 |
| 9 | 4.23 | 3.22 | 2.13 |
| 10 | 4.49 | 3.49 | 2.86 |
| 11 | 4.14 | 2.99 | 2.33 |
| 12 | 4.66 | 3.10 | 2.09 |
| 13 | 3.98 | 2.04 | 1.99 |
| 14 | 4.37 | 2.34 | 2.34 |
| 15 | 4.12 | 2.83 | 2.43 |
| 16 | 4.00 | 3.16 | 2.66 |
| 17 | 4.00 | 3.93 | 3.77 |
| 18 | 3.77 | 3.56 | 3.41 |
| 19 | 3.49 | 3.42 | 3.30 |
| 20 | 3.01 | 2.99 | 2.78 |
| 21 | 2.12 | 2.00 | 1.90 |
| 22 | 2.23 | 1.89 | 1.89 |

**Figure 8:**
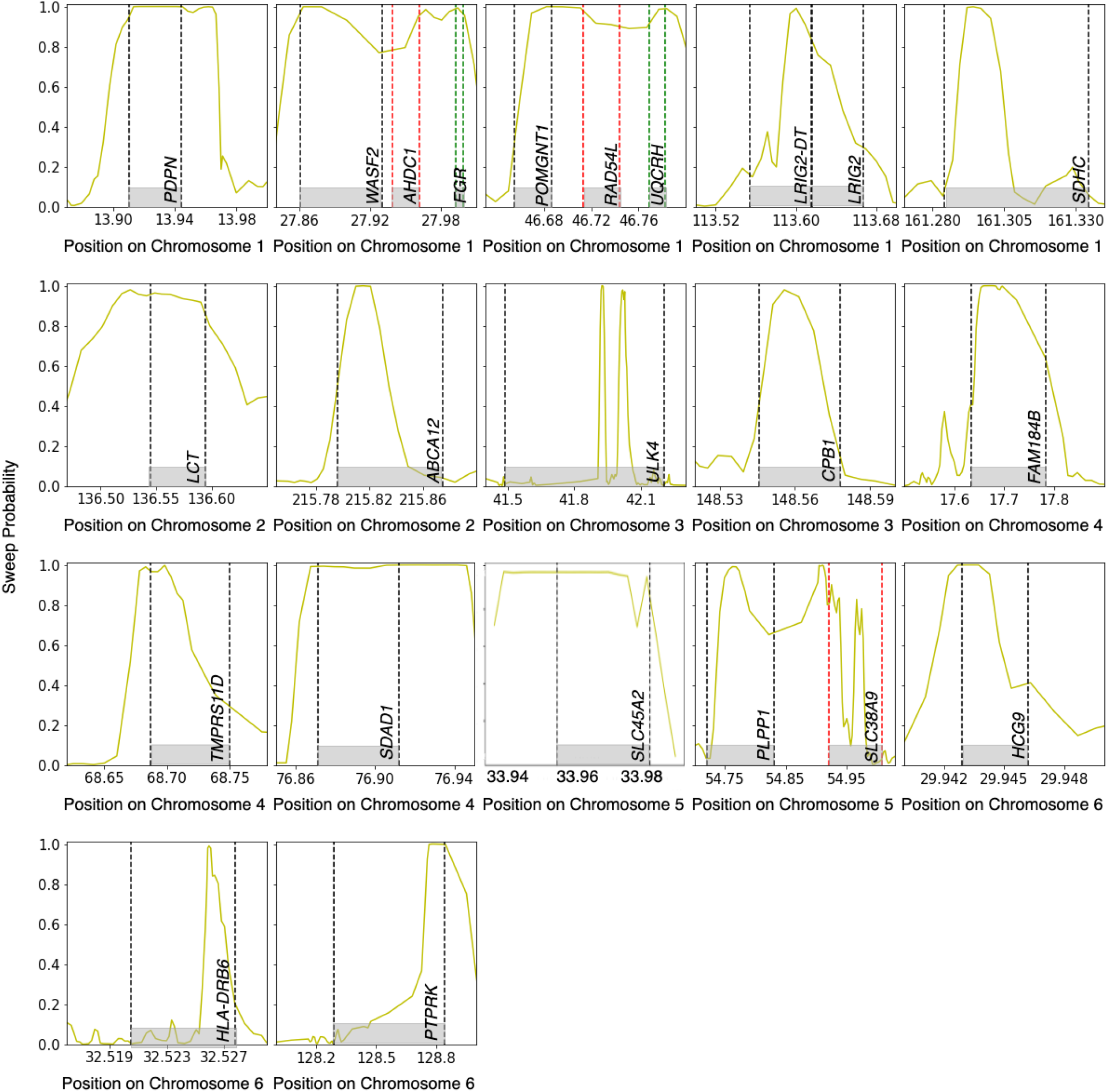
The genome-wide sweep scan results generated by the trained *SISSSCO*[*27CD*] model on the central European humans (CEU population in the 1000 Genomes Project). Ten consecutive windows of sweep probability higher than 0.9 was chosen as the qualifying criteria to be classified as a region to be under positive natural selection. In total 23 genes in 17 regions in the genome show qualifying signs of sweep.

**Table 2:** List of peaks and corresponding genes detected by the *SISSSCO*[*27CD*] model meeting the sweep qualifying criteria of CEU individuals from the 1000 Genomes Project dataset.

| Chromosome | Start (Mb) | Stop (Mb) | Genes |
| --- | --- | --- | --- |
| 1 | 13.83 | 13.97 | *PDPN* |
| 1 | 27.82 | 27.89 | *WASF2* |
| 1 | 27.96 | 28.02 | *ADHC1* |
| 1 | 46.67 | 46.79 | *POMGNT1, RAD54L, UQCRH* |
| 1 | 113.61 | 113.63 | *LRIG2-DT, LRIG2* |
| 1 | 161.29 | 161.31 | *SDHC* |
| 2 | 136.47 | 136.58 | *LCT* |
| 2 | 215.81 | 215.88 | *ABCA12* |
| 3 | 41.88 | 41.94 | *ULK4* |
| 3 | 41.96 | 42.05 | *ULK4* |
| 3 | 148.55 | 148.57 | *CPB1* |
| 4 | 17.61 | 17.74 | *FAM184B* |
| 4 | 68.66 | 66.71 | *TMPRS11D* |
| 4 | 76.83 | 76.93 | *SDAD1* |
| 5 | 33.92 | 33.99 | *SLC45A2* |
| 5 | 54.75 | 54.80 | *PLPP1* |
| 5 | 54.89 | 54.90 | *SLC38A9* |
| 6 | 29.94 | 29.95 | *HCG9* |
| 6 | 32.53 | 32.53 | *HLA-DRB6* |
| 6 | 128.55 | 128.99 | *PTPRK* |

With a predicted sweep probability of 1.0 and a 10-window mean of 0.9998, the *LCT* gene harbors one of the clearest indicators of a sweep found by *SISSSCO*. This high sweep support reinforces the overwhelming evidence for recent positive selection at *LCT* in Europeans from prior studies [*e.g*., Tishkoff et al., 2007, Field et al., 2016, Ségurel and Bon, 2017]. Because of various polymorphisms in the *LCT* gene, which encodes lactase-phlorizin hydrolase, the percentage of adults who are able to digest lactose varies substantially across the world’s populations [Boll et al., 1991]. In particular, the geographical distribution of dairy production and lactase persistence are correlated with one another [Boll et al., 1991]. Moreover, groups where milk and milk products are consumed have been shown to have higher *LCT* gene expression levels [Tishkoff et al., 2007]. High incidence of lactase persistence in European adult populations are the product of positive selection brought about as it prevented lactose intolerance for the people in populations who were consuming dairy products [Bayless and Rosensweig, 1966, Scrimshaw and Murray, 1988]. The *SISSSCO* model suggests that the high-frequency haplotype at *LCT* is the result of one of the most significant recent signals of positive selection in the genomes of Europeans.

Another region that showed high sweep prediction probabilities, including a peak of 0.987 and a 10-window mean of 0.967, is the region containing the *ABCA12* gene, which codes for the protein ATP-binding cassette transporter [Annilo et al., 2002]. The *ABCA12* gene is an absolute requirement for the outer layer of the skin to be able to transport lipids and enzymes [Akiyama, 2014]. This molecular movement is the only way to keep the lipid layers in the epidermis, which are vital to the maintenance of proper skin development [Akiyama, 2014]. The lipid barrier of the skin is the first line of defense that the body has against potentially harmful environmental toxins. Multiple variations of hair and skin pigmentation exist to adapt to different levels of ultraviolet radiation [Baroni et al., 2012, Jablonski and Chaplin, 2010]. A genome-wide scan in Eurasians found that a variant in the *ABCA12* gene harbors footprints of positive selection [Sirica et al., 2019], and *SISSSCO* lends support to these claims with high confidence of a predicted sweep in this region.

Furthermore, the region including the gene *SLC45A2* passed the sweep qualification criterion, with a peak of 0.996 predicted sweep probability and a 10-window mean of 0.9906. The protein coded by *SLC45A2*, which is found in melonocytes, is a key component of the operations responsible for transporting and processing pigmentation enzymes throughout the cell [Kamaraj and Purohit, 2014]. The frequency of an allele in *SLC45A2*, which induces lighter skin pigmentation in modern humans, seems to increase from southern to northern Europe [Costin et al., 2003]. In populations with lighter skin pigmentation, there is a considerable association between regional diversity in multiple functional skin pigmentation polymorphisms within the gene and distance from the equator [Wilde et al., 2014]. This correlation suggests that selection pressure occurred within populations residing in high latitude regions compared to the ones living in lower latitudes over the course of human evolution, as vitamin D3 photosynthesis in northern Europe is expected to be higher for lighter than for darker skin [Novembre and Di Rienzo, 2009, Wilde et al., 2014]. Along with *ABCA12*, the detection of *SLC45A2* by *SISSSCO* lends support to the hypothesis that multiple genes responsible for skin pigmentation went through positive selection in Europeans [Jablonski and Chaplin, 2017].

*SISSSCO* also identified four candidate genes in the major histocompatibility complex (MHC) region. Among them, *HLA-DRB6* and *HCG9* passed our sweep qualification criterion with peaks of 0.9812 and 1.0 predicted sweep probability, and 10-window means of 0.977 and 0.992, respectively. However, the other candidates (*HLA-DRA* and *HLA-A*) moderately exhibit signatures of sweeps, as they do not pass the stringent qualification criterion, but do pass it if we relax the threshold to a 10-window mean of 0.7. Though categorized as a MHC gene, *HLA-DRB6* is a pseudogene [Cree et al., 2010] that may have lost its first exon and promoter to the insertion of a virus far in the past, thereby making it nonfunctional [Mayer et al., 1993]. In contrast, *HCG9* is a long non-coding RNA gene [Pal et al., 2016], and hence may be involved in gene regulation. The MHC region contains many exceptionally highly polymorphic genes that code for cell surface proteins responsible for communication between cells and extracellular environments [Horton et al., 2004]. These proteins make up the adaptive immune system by recognizing foreign pathogens to initiate a targeted immune response, which becomes essential when the innate immune system fails in protecting cells [Horton et al., 2004]. Among MHC class I genes, *HLA-A* showed marginal signs of positive selection with 10-window mean sweep prediction probability of 0.70. Similarly, among MHC class II genes, along with *HLA-DRB6*, *HLA-DRA* showed signs of positive selection with 10-window mean sweep prediction probability of 0.72. The marginal sweep candidates *HLA-A* and *HLA-DRA* show a trend of multiple genes in the MHC class I and class II to exhibit signs of sweeps. These findings are reinforced by other studies that observed sweep signatures at the MHC region within Europeans [*e.g*., Albrechtsen et al., 2010, Goeury et al., 2018, Harris and DeGiorgio, 2020b, DeGiorgio and Szpiech, 2022].

*SISSSCO* detected 16 other sweep candidates, a large number of which are associated with cancer detection or suppression. Specifically, the *PDPN* gene that encodes the protein Podoplanin, which serves as a marker for lymphatic vessels [Kitano et al., 2010]. Because it can be utilized as a tool, though rather weak, for cancer diagnosis, this gene has played a crucial role in cancer research [Kawaguchi et al., 2008, Krishnan et al., 2018, Quintanilla et al., 2019]. Additionally, it is a major factor in the metastasis of squamous cell carcinoma, a common form of skin cancer [Kitano et al., 2010]. The genes *WASF2* and *LRIG2* have been linked with many forms of cancer detection as well [Wang et al., 2014, Kitagawa et al., 2019]. *WASF2* expression levels have been studied as a biomarker in detection of pancreatic [Kitagawa et al., 2019] and ovarian cancers [Yang et al., 2022], whereas *LRIG2* has been identified as a biomarker for detection of non-small cell cancer [Wang et al., 2014]. On the other hand, *SDAD1* has been identified as a gene responsible for suppressing colon cancer metastasis [Zeng et al., 2017]. A number of prior scans also found traces of selective sweep footprints in cancer associated genes. For instance, Lou et al. [2014] and Mughal and DeGiorgio [2019] identified the *BRCA1* gene as a sweep candidate, and Schrider and Kern [2017] detected sweep signatures at several cancer related genes, including *CADM1* and *MUPP1*. Though the cancer associated genes detected by *SISSSCO* differ from those of prior studies, these findings portray an interesting enough trend that *SISSSCO*, along with a number of other approaches from prior studies, identified several cancer related genes as selective sweep candidates.

## Discussion

In this study, we found that the *SISSSCO* models do indeed have increased power and accuracy compared to the three competing summary statistic-based machine learning methods. In particular, the 27-input CNN models (*SISSSCO*[*27CO*], *SISSSCO*[*27CD*], and *SISSSCO*[*27MD*]) generally outperformed the three-input CNN models (*SISSSCO*[*3CO*], *SISSSCO*[*3CD*], and *SISSSCO*[*3MD*]), with all three 27-input models showing similarly high performance across tested demographic histories and selection regimes. Though classification accuracy is slightly lower for *SISSSCO*[*27CD*] than for *SISSSCO*[*27MD*] on missing data, given its high accuracy and power across the range of demographic and adaptive scenarios tested as well as robustness to missing data, we decided to use this method to detect sweeps on an empirical human genomic dataset.

Application of *SISSSCO* to European human genome variation gave high support for previously-identified sweeps at the *LCT*, *ABCA12*, and *SLC45A2* genes [Bersaglieri et al., 2004, Beleza et al., 2013, Sirica et al., 2019], as well as 19 other candidate genes with high confidence. We employed a stringent sweep qualification criterion to limit the number of falsely discovered sweeps. A key finding is that, two genes in the MHC region, namely *HCG9* and *HLA-DRB6*, and with a relaxed qualification criterion another two genes *HLA-A* and *HLA-DRA*, presented sweep signatures. However, past studies have indicated that the MHC region has undergone balancing selection [*e.g*., Solberg et al., 2008, Cagliani et al., 2011]. As recent balancing selection leaves a spatial pattern in the genome similar to that of an ongoing selective sweep [Isildak et al., 2021], *SISSSCO* may have picked up such spatial patterns in the MHC region. Moreover, we found that, as expected, the vast majority of the genomic windows were classified as neutral, with only a handful of regions showing clear sweep signatures. These results echo those from simulated data that *SISSSCO*[*27CD*] has high accuracy to discriminate sweeps from neutrality, while remaining robust to false discovery due to technical artifacts expected in real empirical datasets.

The three spectral analysis techniques that we employed add versatility to *SISSSCO*, as they focus on different characteristics of signals. In particular, they extract information from multiresolution analysis of low- and high-frequency regions within the summary statistic signals. This information is obtained either through wavelet transformation of signals or through multitaper spectral analysis by tapering signals using qualifying tapers to generate power maps emphasizing overall signal shapes. Focusing on genomic spatial windows as a function of the dominant frequency within the summary statistic signal through the S-transform also offers a unique mechanism for drawing information from signals. By leveraging these diverse patterns of information, *SISSSCO* gains the ability to build a more accurate and robust system compared to existing sweep detectors that utilize vectors of multiple summary statistics as input.

To study the benefits of adding the layer of spectral inference within *SISSSCO*, we evaluated the accuracy and power of CNN models that take as input nine raw summary statistic vectors instead of 27 time-frequency analysis images. Specifically, we adapted the *SISSSCO* model architectures to construct four one-dimensional CNN models: a single CNN with nine channels (*1D-CNN*[*1CNN*]), nine pretrained single-channel CNNs with the output layers concatenated (*1D-CNN*[*9CO*]), nine pretrained single-channel CNNs with the dense layer concatenated (*1D-CNN*[*9CD*]), and nine simultaneously-trained single-channel CNNs with the dense layer concatenated (*1D-CNN*[*9MD*]). We find that all four *1D-CNN* methods have substantially lower classification accuracy and power than *SISSSCO*[*27CD*] on the `Nonequilibrium_variable` dataset (compare Figure S16 to Figure 4). Among the four *1D-CNN* models, we found *1D-CNN*[*9MD*] to have highest accuracy, which is approximately five percent lower than *SISSSCO*[*27CD*]. The powers of the *1D-CNN* methods evidenced by the ROC curves echo the relative accuracies of the methods, with the ranking from worst to best performance given by *1D-CNN*[*1CNN*], *1D-CNN*[*9CO*], *1D-CNN*[*9CD*], and *1D-CNN*[*9MD*]. The powers demonstrated by the *1D-CNN* architectures are dwarfed by *SISSSCO*[*27CD*], which displays a near perfect area under the ROC curve (Figure S16). Though the *SISSSCO* models require significantly more time and computational resources to train compared to the *1D-CNN* models, the improvement in model performance is quite considerable. Therefore, adding the layer of spectral inference appears to provide additional performance gains to *SISSSCO* compared to operating on the raw summary statistics.

A potential reason that these signal decomposition methods offer improved predictive ability over the use of raw summary statistics might be that they aim to isolate low-frequency components that are responsible for overall trends of signals, but place lower importance on regions of signals where abrupt changes occur. These low-frequency components may be generated by genetic variation within the population stemming from nonadaptive processes including mutation, recombination, migration, and genetic drift. However, the regions of abrupt changes on top of the scope of the underlying low-frequency components may be caused by adaptive processes, such as positive natural selection.

The two-dimensional images generated by these time-frequency analysis tools have different patterns that can be used to explain the energy of the high-frequency components within the signal. CNNs therefore play a vital role in identifying regions of interest from these images. Because CNNs are so flexible, we were able to set up image processing architectures that were suited for finding specific regions of interest in the three types of images made by the three signal decomposition methods that match the complexity of patterns in those regions. In addition to this adaptability, the CNNs made it possible to combine data from several image types to create a stacked (or concatenated) set of models with increased ability to spot signs of adaptive events.

We tested six stacking, or concatenation, architectures that utilize information from 27 input images generated by nine summary statistic signals, each decomposed with three time-frequency analysis methods. Three of our six stacked models involve three nine-channel input CNNs (*SISSSCO*[*3CO*], *SISSSCO*[*3CD*], and *SISSSCO*[*3MD*]), whereas the other three operate on 27 single-channel input CNNs (*SISSSCO*[*27CO*], *SISSSCO*[*27CD*], and *SISSSCO*[*27MD*]). The three stacking approaches involving nine-channel input CNNs generally performed better than each of the signal decomposition methods tested in isolation as presented in the Results section, corroborating the motivation that combining knowledge from three signal decomposition methods does indeed enhance classification performance. A likely reasoning for this result is that the different signal decomposition methods interrogate distinct properties of a signal, making images from the three time-frequency analysis approaches complementary rather than redundant. On the other hand, stacking methods employing nine-channel CNNs were often outperformed by those employing single-channel CNNs.

Thus far, we have focused on the predictive ability of the *SISSSCO* models. However, interpretability of the models is also important. A mechanism that can facilitate interpretation is through computation of saliency and class activation maps [Zhai and Shah, 2006]. When discussing visual processing, the term “saliency” refers to the ability to recognize and differentiate individual aspects of an image, such as its pixels and resolution. These elements highlight the most visually compelling parts of an image. Saliency maps are a topographical representation of these locations. The goal of saliency maps are to reflect the degree of importance of a pixel to the human visual system. Computing mean saliency and class activation maps might lend us the opportunity to intuitively understand the CNN models implemented in *SISSSCO*, by permitting visualization of the particular regions of images that trained CNNs place emphasis for detecting adaptation. It may also provide us with the information as to how the single-channel CNNs outperformed the nine-channel CNNs. Incorporation of class activation and saliency maps through attention has been demonstrated to be effective in recent deep learning methods for detecting adaptive introgression [Gower et al., 2021], and our models could be adjusted to accommodate a similar approach.

Across the various simulated test settings, the relative performances of the *SISSSCO* and non-*SISSSCO* models remained consistent, as did the relative performances among non-*SISSSCO* models. A comprehensive understanding of what drives these differences in classification behavior is difficult, but key characteristics of modeling decisions may provide some light. First, though *SISSSCO* and *SURFDAWave* both employ signal decomposition methods as well as the same set of summary statistic vectors, the underlying relationships between the class labels and the summary statistic values may be nonlinear, and thus the nonlinear CNN models employed by *SISSSCO* may provide it with better accuracy and power. Moreover, three signal decomposition methods employed by *SISSSCO* each interrogate different characteristics of a signal and are thus complementary. In contrast *SURFDAWave* considers only a single signal decomposition method for extracting features from summary statistic vectors.

Next, diploS/HIC uses a different set of summary statistics that operate on unphased multilocus genotype data. This use of unphased multilocus genotype summary statistics may be why diploS/HIC has lower accuracy and power than *SURFDAWave* and some of the *SISSSCO* models on the evolutionary settings tested, as using input summaries computed from unphased rather than phased data could decrease classification accuracy [Mughal et al., 2020]. Second, diploS/HIC divides the analyzed genomic region into a small number of large physical-based windows, whereas *SISSSCO* uses a large number of SNP-based windows. These SNP-based windows give *SISSSCO* robustness to missing genomic regions, whereas diploS/HIC is less robust due to its use of physical-based windows—though masking of genomic regions can be implemented within model training to account for missing data [Kern and Schrider, 2018]. Third, diploS/HIC does not use ensemble learning other than dropout layers. Fourth, diploS/HIC normalizes each summary statistic across windows, whereas *SISSSCO* does not normalize summary statistic signals before signal decomposition. Finally, diploS/HIC was designed to discriminate among five classes, which is important because the diploS/HIC summary statistics may have been chosen to provide optimal performance for the original setting of five classes.

In relation to evolBoosting, though it employs ensemble learning similar to *SISSSCO*, these ensemble approaches have many differences. That is, evolBoosting utilizes boosting, which aggregates predictions from many weak learners, whereas the stacking approach of *SISSSCO* takes node weights from the fully connected dense layers or the output layers of the individually trained CNNs, which are each potentially strong predictive models. Second, evolBoosting uses a different set of summary statistics, computed across a moderate number of moderate-length physical-based windows. Similarly to diploS/HIC, this sensitivity of evolBoosting to missing data is likely due to the calculation of summary statistics of physical-based windows.

The *SISSSCO* models were trained with phased haplotypic data. However, phased data is difficult or impossible to reliably generate for many study systems—notably most nonmodel organisms. Hence, for our models to be versatile, it is imperative that they can also accommodate unphased data (*e.g*., similarly to diploS/HIC of Kern and Schrider [2018]). Fortunately, the phased haplotype summary statistics used by *SISSSCO* have natural analogs for unphased multilocus genotype data. Specifically, we could replace *H*_1_, *H*_2_/*H*_1_ and *H*_12_ with their respective unphased analogs *G*_1_, *G*_2_/*G*_1_, and *G*_123_ [Harris and DeGiorgio, 2020a] and exchange the frequencies of the five most common haplotypes with the five most common unphased multilocus genotypes. Given the relatively strong concordance with results from haplotype-based methods [Harris and DeGiorgio, 2020a, DeGiorgio and Szpiech, 2022, Harris and DeGiorgio, 2020b] and power to detect sweeps in prior studies using unphased multilocus genotypes [Kern and Schrider, 2018, Mughal and DeGiorgio, 2019, Gower et al., 2021], we expect that *SISSSCO* would retain excellent classification accuracy and power when applied to unphased data.

Though we focused on the application of *SISSSCO* to binary classification problems, it can be easily extended to multiclass problems and retooled to predict evolutionary parameters within a regression framework, which can provide a richer understanding of the processes that have led to selection footprints in the genome. For example, estimating the timing (*t*) and strength (*s*) of selection may provide a hint at the environmental pressures that led to the rise in frequency of particular traits associated with identified sweep candidates. Moreover, predicting the frequency of the allele when it became adaptive (*f*) can lend information about the mode of positive selection at candidate genes, with low frequency suggesting a hard sweep from a *de novo* mutation and moderate frequency a soft sweep from standing variation. Furthermore, incorporating images of twodimensional statistics may be helpful when considering multiclass problems, such as discriminating among neutrality, non-introgression sweeps, and adaptive introgression [Racimo et al., 2015]. In particular, Mughal et al. [2020] showed that including two-dimensional statistics (*i.e* moments of the distribution of the squared correlation coefficient *r*^2^ [Hill and Robertson, 1968]) in addition to one-dimensional statistics can aid in discriminating among different types of adaptive processes, such as adaptive introgression and non-introgression sweeps. However such two-dimensional summary statistics do not fall within the *SISSSCO* framework developed here. Instead, *SISSSCO* could accommodate images that are not from spectral analysis, such as moments of pairwise linkage disequilibrium computations, as separate concatenation branches.

Overall, spectral analysis of genomic summary statistics that result in time-frequency images offer precise localization of high-frequency components within the signal. In contrast to the frequency components generated by genetic variation due to nonadaptive events, the high-frequency components caused by adaptive events like positive natural selection are qualitatively different. This article also demonstrated that stacking is a useful technique for integrating models that search for signatures of such evolutionary events in various ways. We further believe that *SISSSCO* will prove to be a powerful tool for future development of robust predictive models that aim to find traces of adaptive events, and predicting evolutionary parameters by tapping into the potential of spectral analysis.

## Methods

### Computing *SISSSCO* summary statistics from simulated data

For the purpose of training the *SISSSCO* models, we generated the nine summary statistics from the population sample files that we simulated using `discoal`. As discussed in the *Modeling description* subsection of the *Results*, we generated four simulated training sets: `Equilibrium_fixed`, `Equilibrium_variable`, `Nonequilibrium_fixed`, and `Nonequilibrium_variable`. We parsed each replicate from these simulated datasets to include the central 400 SNPs (200 to the left of the sequence center and 200 to the right of the sequence center). Using these 400 SNPs, we calculated the nine summary statistics for our training, validation, and test sets with a window of size 10 SNPs and a stride of three SNPs. This procedure resulted in summary statistic arrays of length 128 windows. The nine summary statistic vectors of size 128 were then fed into the three signal decomposition methods with identical protocols and packages [Cokelaer and Hasch, 2017, Satriano, 2017, Lee et al., 2019] as described in the *Modeling description* subsection of the *Results*. As a result, a total of 27 time-frequency analysis images of size 65 × 128 were generated per simulated replicate.

### Spectral analysis of summary statistics

Each of the nine summary statistics described in this study exhibit oscillatory dynamics. The oscillatory characterization of time series data provides valuable insights into the construction of the data via spectral analysis [Babadi and Brown, 2014]. However, for our purpose, we calculated these summary statistics over contiguous windows, which portray autocorrelation properties similar to that of time series data. A key characteristic of our summary statistic computations is that they are of finite length, while in theory we need a sample of infinite length to describe a system in the frequency domain. However, finite-length data can result in spectral analysis that is highly erroneous [Sadowsky, 1996, Babadi and Brown, 2014]. In this subsection, we consider three different methods for performing spectral analysis on finite-length signals: wavelet decomposition, multitaper analysis, and the S-transform. Furthermore, we generally follow the notation of Sadowsky [1996], Babadi and Brown [2014], and Yun et al. [2013] to respectively describe the wavelet decomposition, multitaper analysis, and S-transform, with modifications to ensure uniform and consistent notation across subsections.

#### Wavelet decomposition

The continuous wavelet transform (CWT) permits the examination of signals, the extraction of spectral features, and the discovery of nonstationary properties that are dependent on time and scale [Sadowsky, 1996]. It is a technique that takes a signal *x*(*t*) over time *t* and produces a time- and scale-variable parameter surface that could prove useful for its characterization of a signal and the origin of the signal. For the CWT to fulfill the requirements of its role as the kernel function of a signal transform, it is specified in relation to a basis function *ψ*(*t*) termed a mother wavelet. To qualify as a mother wavelet, a wavelet must satisfy two properties. The first property is that the mother wavelet is designed so that the wavelet transform is invertible [Sadowsky, 1996]. That is, because the wavelet transform takes a signal from the time domain and projects it onto a timefrequency plane, there must be an operation that permits the reconstruction of the time domain signal from the time-frequency plane. In addition to this property, the “admissibility condition” must also be met by the mother wavelet. The admissibility condition states that, for there to be an inverse wavelet transform, the Fourier transform [Grafakos, 2008] of the mother wavelet must be zero for any constant component in the signal, and thus have zero direct current bias [Holschneider, 1996]. Therefore, the mother wavelet must have oscillations to meet the admissibility condition [Sadowsky, 1996].

The Fourier transform is a mathematical tool used for frequency analysis of a signal, which transforms a time domain signal into the frequency domain. That is, it is a method of frequency domain representation of a signal, which can also be reversed to get the time domain signal. The Fourier transform employs a technique so that every signal can be decomposed into one or many sinusoidal waves of varying frequencies and amplitudes. For a continuous time signal *x*(*t*), the transformation is defined as [Bracewell and Bracewell, 1986]

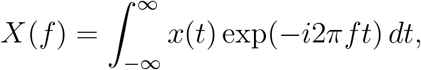

where 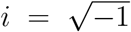 indicates an imaginary component, *f* is the frequency, and complex number exp(−*i*2*πft*) = cos(2*πft*) + *i*sin(2*πft*) can be broken into cosine and sine functions. The real valued waveform cos(2*πft*) and imaginary valued waveform sin(2*πft*) are of same frequency *f*. The product of exp(−*i*2*πft*) with the time domain signal *x*(*t*) gives us the amplitude of every participating waveform in the frequency space.

In this study, we consider the Morlet wavelet as the mother wavelet [Kronland-Martinet et al., 1987]. The Morlet wavelet can be defined as [Sadowsky, 1996]

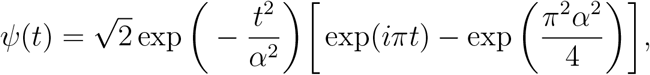

where *α* denotes the shaping factor to obtain a desired time-frequency shaping of the Morlet wavelet. Time-frequency shaping helps generate a time-frequency image with resolution and size that is suitable for a given performed analysis. The frequency domain representation of the mother wavelet after applying the Fourier transform is

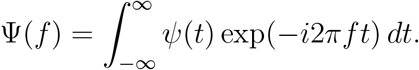

Because the admissibility condition dictates that *ψ*(0) = 0, it follows that [Sadowsky, 1996]

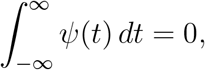

which leads to the frequency domain representation of the Morlet wavelet as

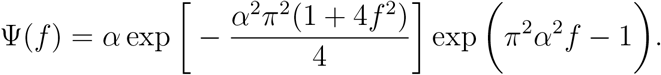

We set the shaping factor as 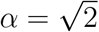 to ensure reduction of frequency overlap while preserving a reasonable level of temporal resolution. This a value results in horizontal shaping of the mother wavelet in the time domain to obtain the necessary number of oscillations (Figure 9), and determination of the center frequency of the wavelet in the frequency domain.

**Figure 9:**
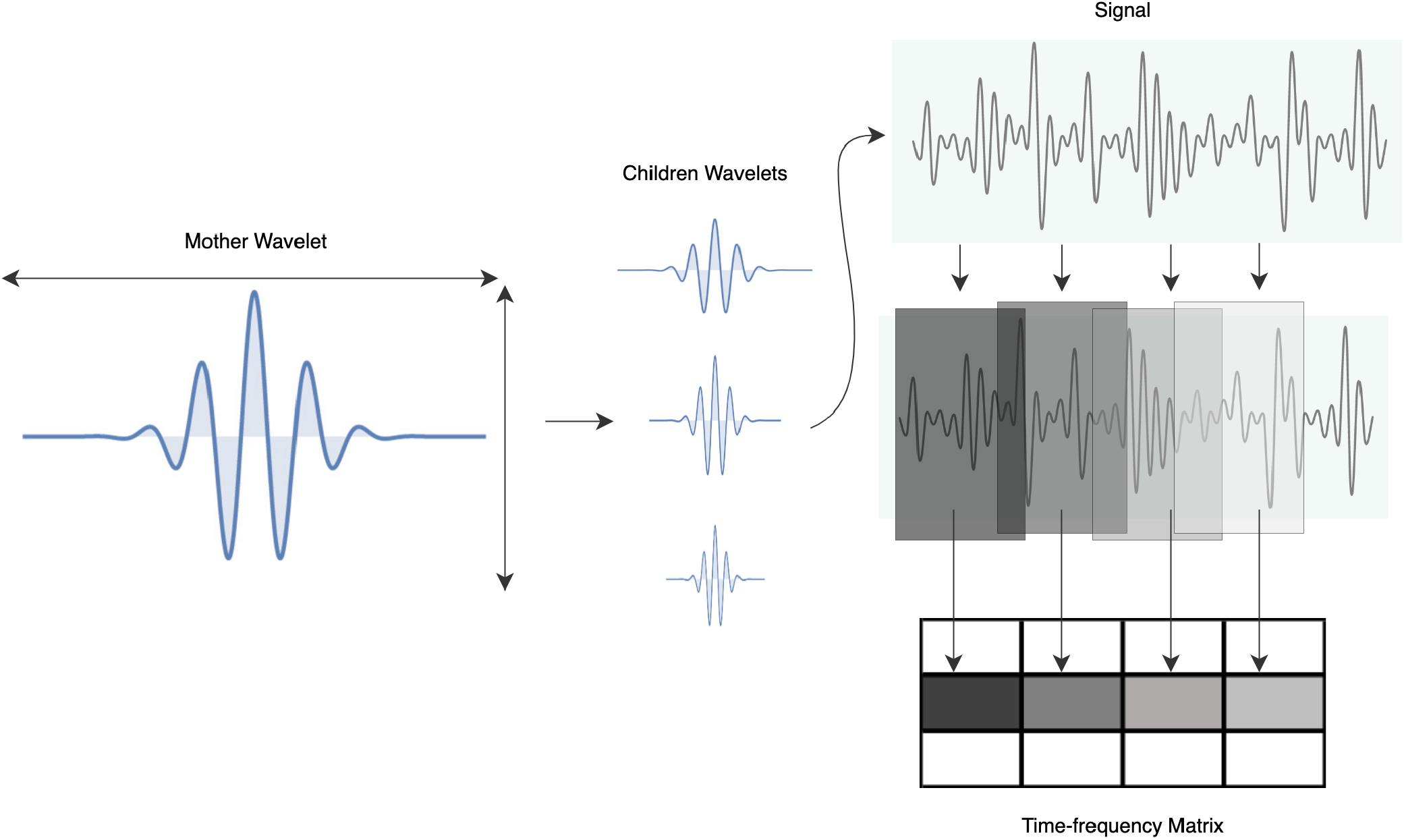
Process flow of generating the time-frequency analysis matrix using wavelet transform. The “mother wavelet” is stretched and squeezed both horizontally and vertically to produce different realizations of the wavelet having different scales known as “children wavelets”. Each of these children wavelet are translated or time-shifted across the signal to match the wavelet against continuous windows of the signal to get the match coefficients that generate the individual rows of the time-frequency analysis matrix. Higher match results in higher coefficient values, whereas lower match results in lower coefficient values.

The CWT of a signal *x*(*t*) with respect to a wavelet *ψ*(*t*) is a function of scaling factor *a* and translation factor *b*, and can be expressed as [Daubechies, 1992, Sadowsky, 1996]

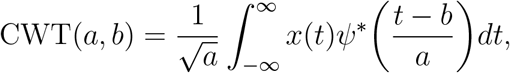

where the superscript * indicates complex conjugation and where locality in time and frequency are controlled by parameters *b* and *a*, respectively. Scaling can refer to either a reduction or an increase in horizontal shape, as it can be both contracted (squeezed) or dilated (stretched). It is feasible to express the amplitude versus the scale and its fluctuation over time by altering the scale and translation parameters along the time index *t*. The wavelet is said to be stretched if *a* > 1, and squeezed if 0 < *a* < 1. In this study, the translation parameter is discretized to integer values, whereas the scale parameter is discretized to fractional powers of two.

Figure 9 depicts the mother wavelet and its children wavelets produced by changing scale factors. Fixing the scaling factor *a*, we perform the CWT(*a, b*) with increasing values of translation factor *b*. The translation, represented by the shaded blocks in Figure 9, makes up each row of the multiresolution time-frequency image, which is referred to as a scalogram.

#### Multitaper analysis

Multitaper analysis is a nonparametric method introduced to overcome the high bias and error variance of time series data [Berardi and Zhang, 2003]. Bias is the discrepancy between the expected value of an estimator and the true underlying function, whereas variance refers to the spread of the distribution of functions about this expected value [Berardi and Zhang, 2003]. Multitaper analysis attempts to overcome a key limitation of conventional Fourier analysis, as it does not assume that a single instance of a noisy statistical signal can deliver the true coefficients of the underlying process of interest [Prieto et al., 2007]. To decompose a signal into one or many sinusoidal waves of varying frequencies and amplitudes, the Fourier transform assumes that the signal is of infinite length. However, the summary statistic vectors that we employ as our signals are of finite length.

Using the frequency analysis of a time series that has been discretized over time, the Fourier transform can be expressed as [Bracewell and Bracewell, 1986]

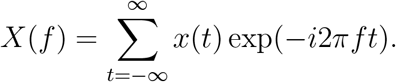

Let *x_k_* = *x*(*k*Δ), *k* = 0, 1, …, *N* – 1, be a discrete-time signal of finite length *N* for sampling interval Δ. That is, the underlying continuous analog signal *x*(*t*) from which the finite-length digital signal *x_k_* is generated was sampled after every Δ time unit. The Fourier transform of *x_k_* is defined as [Babadi and Brown, 2014]

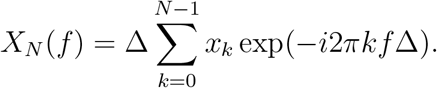

The power spectral density [Youngworth et al., 2005] defines the distribution of a signal’s power as a function of frequency *f* and aids in the identification of the frequency ranges where changes in the signal are prominent. To compute the power spectral density, the mean power *P*(*f*) in the frequency band of 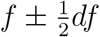, where *df* indicates an arbitrarily small amount of change in frequency *f*, is defined as [Babadi and Brown, 2014]

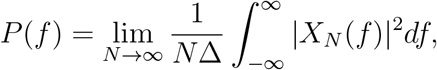

where *X_N_*(*f*) is the frequency domain representation of *x_k_*, *k* = 0, 1, …, *N* – 1, and the expression 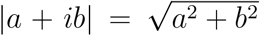 denotes modulus of complex number *a* + *ib*. However, because as *N* approaches infinity there are never enough windows *N* in real-world settings, it is impossible to compute this quantity in practice. Instead, we constrain the analysis to second-order stationary and ergodic sequences, as the summary statistic vectors in this study are computed from a finite number of genomic windows. A constant mean and a time-invariant autocovariance are two crucial characteristics of second-order stationary signals [Boshnakov, 2011]. On the other hand, any given reasonably sized sample from an ergodic process can be taken as a true reflection of the process [Cherstvy et al., 2013].

According to the Wiener-Khintchine theorem [Khintchine, 1934], the power spectrum of a wide-sense stationary random process, such as a second-order stationary process, can be used to derive the spectral decomposition of the autocovariance function *s_k_, k* = 0,1, …, *N* – 1, of the process, with *s_k_* = 0 for all other values of *k*. This theorem dictates that

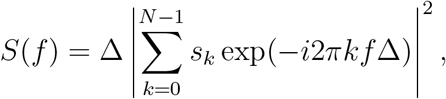

where *S*(*f*) is the power spectral density of the discrete window signal at frequency *f*. Computing of an ergodic second-order stationary infinite length signal *x_k_*, –∞ < *k* < ∞, would provide the power spectral density [Babadi and Brown, 2014]. However, we do not have an infinite-length signal. Assume that 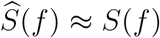, where 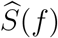 is the power spectral density estimated from finite-length signal *x_k_*, where the variance of the estimated power spectral density is approximately zero. Denoting the autocovariance of *x_k_* by 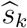, *k* = 0,1, …, *N* – 1, the Fourier transform of the sequence 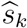 yields the power spectral density [Bartlett, 1950, Babadi and Brown, 2014]

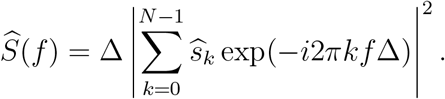

Bias is the distinction between the true power spectral density and a smoothed representation of the true power spectral density, which can be divided, at a given frequency, into narrow-band bias and wide-band bias. The dominant frequency components cause narrow-band bias, whereas the minor ones cause wide-band bias. Consider a taper *h_k_*, *k* = 0,1, …, *N* – 1, which when multiplied with *x_k_*, generates a tapered sequence (see Figure 10). A periodogram estimate of this tapered sequence can be written as [Babadi and Brown, 2014]

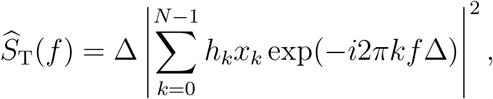

where the signal is replaced by the product of a taper *h_k_* and the signal *x_k_*. Tapering presents a middle ground between narrow-band and wide-band bias that helps equalize the imbalance of these two forms of biases [Bronez, 1992, Babadi and Brown, 2014]. Multitaper spectral estimation is used to distinguish between optimal tapers and suboptimal tapers, which are unable to efficiently localize the frequency components. High variance for *N* ≫ 1 is a drawback shared by both the 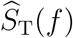 and 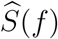 estimates, and this variance does not converge to zero as *N* approaches infinity, preventing these estimates from exactly matching the true power spectral density. Multitaper spectral analysis aids in overcoming this drawback.

**Figure 10:**
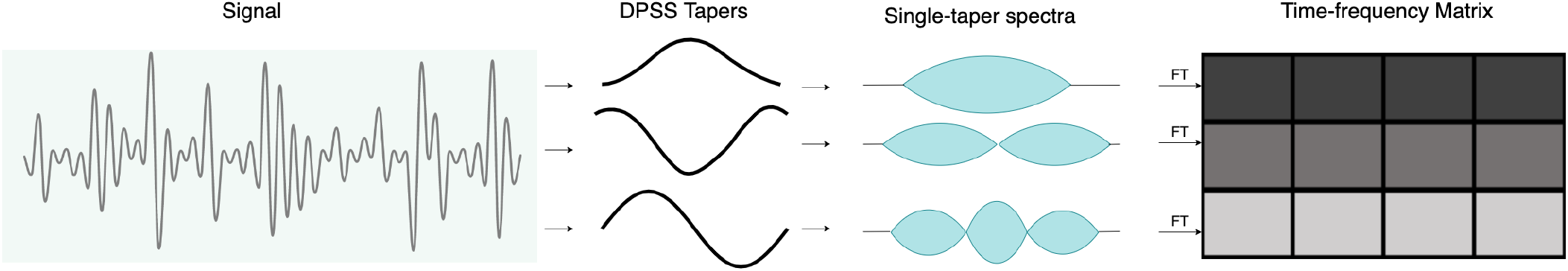
Process flow of generating the time-frequency analysis matrix using multitaper spectral analysis. A fixed number of DPSS tapers are generated which are designed to extract information from certain characteristics of the signal. Each of the tapers are used to taper the signal that produces the expectation of the periodogram, which is the the single taper spectra produced by a single taper. This single taper spectra is fast Fourier transformed to create the individual rows of the time-frequency analysis matrix.

For a set {*h*_*k*0_, *h*_*k*2_, …, *h*_*k*(*L*–1)_} of *L* uncorrelated tapers each with unit variance, the multitaper spectral estimate of the true power spectral density is defined as [Welch, 1967, Babadi and Brown, 2014]

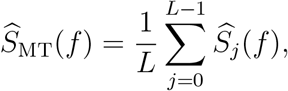

where

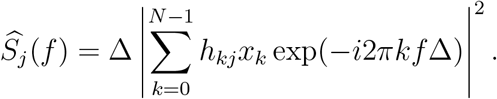

The single-taper spectrum denoted by 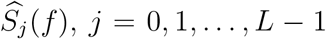, generates each row of the timefrequency matrix (Figure 10). Due to their superior defense against spectral leakage that causes a reduction in frequency resolution [Lyon, 2009], discrete prolate spheroidal sequences (DPSS) [Lees and Park, 1995, Karnik et al., 2022] are often utilized as tapers for the multitaper spectral analysis (Figure 10). Calculating the DPSS tapers that connect frequency resolution to data window size requires the usage of the time-half bandwidth parameter, which is the product of the duration of the data window and half the bandwidth [Prerau et al., 2017].

#### S-transform

Time series characteristics are said to be stationary if they do not change as the series progresses across observational time. Means, variances, and covariances among observations, however, tend to change with time or are non-stationary. In many real-world applications, such as seismographic activity detection and financial forecasting [Frohlich et al., 1982, Abu-Mostafa and Atiya, 1996], it is unrealistic to expect stationarity in a time series, and thus, assuming stationarity may not be particularly useful for characterizing the signal source. Considering the analysis may imply relationships among variables where none exist, drawing a conclusion based on non-stationary time series analysis carries the risk of false interpretation [Stockwell et al., 1996].

Alternately, by utilizing the Fourier transform to convert a signal from the one-dimensional time domain to the one-dimensional frequency domain, we are able to glean further insight into the relationship that exists between the signal *x*(*t*) and its origin (source generating the signal). The signal that is generated as a result of this transformation of domains has a high frequency resolution but a low time resolution. Time-frequency analysis methods involve projecting onedimensional non-stationary signals into a two-dimensional time-frequency plane so that they can be analyzed. To accomplish this projection, the S-transform [Stockwell et al., 1996] makes use of a moving and scalable Gaussian window in conjunction with the concepts behind the short-time Fourier transform [Yun et al., 2013].

Denoting the time-dependent localizing Gaussian window as *w*_G_(*t*), we can write the short-time Fourier transform as [Fano, 1950]

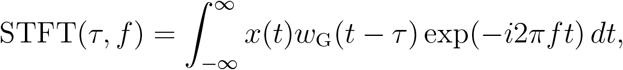

where *τ* is an arbitrary time displacement, and *w*_G_(*t* – *τ*) explains the translational property of the Gaussian window. Stockwell et al. [1996] defines this time-dependent Gaussian window as

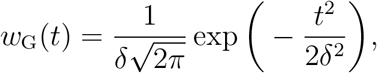

where *δ* is the window width. The horizontal width of the Gaussian window can be adjusted by using the scale factor *δ*. Yun et al. [2013] defines *δ* as *δ*(*f*) = 1/|*f*| so that it is a function of frequency, and thus defines a new time-frequency dependent Gaussian window function as

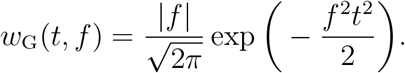

The Gaussian window function with a certain scale factor *δ* is depicted in Figure 11. This window has unit area above the horizontal time axis such that 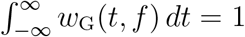, which signifies that the window does not have a diminishing impact on the windowed signal. To expand upon this idea, suppose we have *x*(*t*) = 1 and 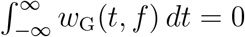, which indicates that the window function has an equal area both above and beneath the time axis. The area under *x*(*t*) is positive. If we multiply *x*(*t*) with *w*_G_(*t, f*) as depicted in Figure 11, then the resultant signal will have an equal area above and beneath the horizontal axis. However, if we have 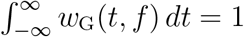, then the resultant signal will also have all of its area above the horizontal axis, which signifies that this Gaussian window preserves the trend of the signal. Putting it all together, the S-transform is defined as [Yun et al., 2013]

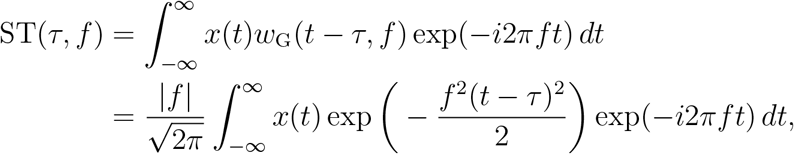

which is the Fourier transformation of the multiplication of the window function and the signal as visualized in Figure 11.

**Figure 11:**
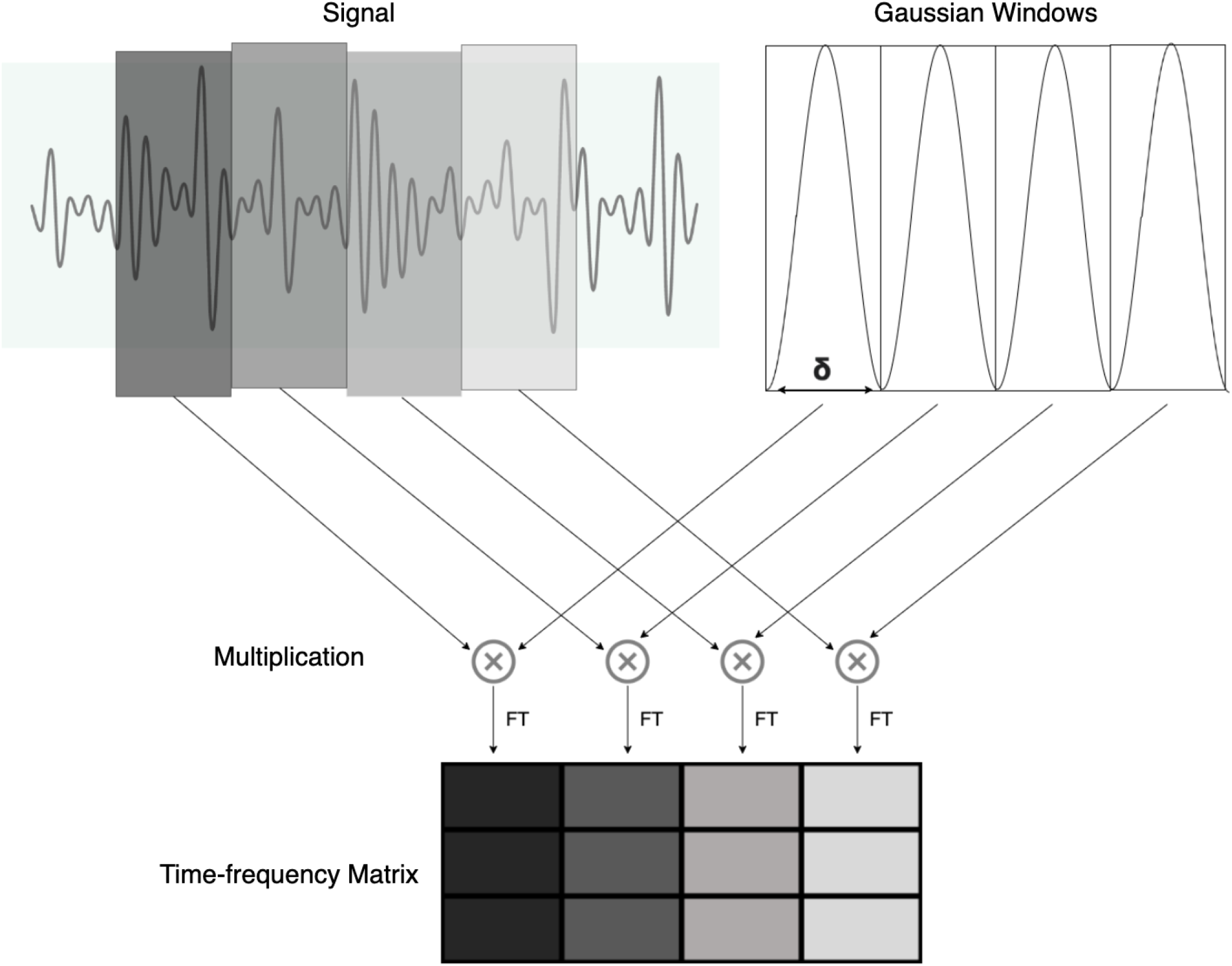
Process flow of generating the time-frequency analysis matrix using S-transform. The signal is multiplied by the Gaussian window function of a certain frequency dependent width at specific non-overlapping intervals that result in a windowed version of the signal. The individual windowed signal is then fast Fourier transformed to produce individual columns of the time-frequency analysis matrix.

### Processing empirical data

Before calculation of the summary statistics from our empirical dataset, we removed the SNPs with a minor allele count of two or lower. To avoid spurious signals due to technical artifacts, we also removed 100 kb regions of mean CRG mappability and alignability score lower than 0.9 [Talkowski et al., 2011]. After the removal of unqualified SNPs, we calculated the nine summary statistics in a identical way to our training dataset, with a window size of 10 SNPs and a stride of three SNPs. To match the length of the summary statistic vectors employed by our trained models, we took 128 consecutive windows of each summary statistic, moving by a stride of one window across each chromosome to generate each additional summary statistic vector until the last window of a particular chromosome is reached. Identical to the process discussed in the *Computing SISSSCO summary statistics from simulated data* subsection, we then generated the 27 time-frequency images from these summary statistic arrays to make our predictions.

## Acknowledgments

This work was supported by National Institutes of Health grant R35GM128590 and by National Science Foundation grants DEB-1949268, BCS-2001063, and DBI-2130666. Computations for this research were performed using the services provided by Research Computing at the Florida Atlantic University.

