## Supplementary Material for "Uncovering footprints of natural selection through time-frequency analysis of genomic summary statistics"

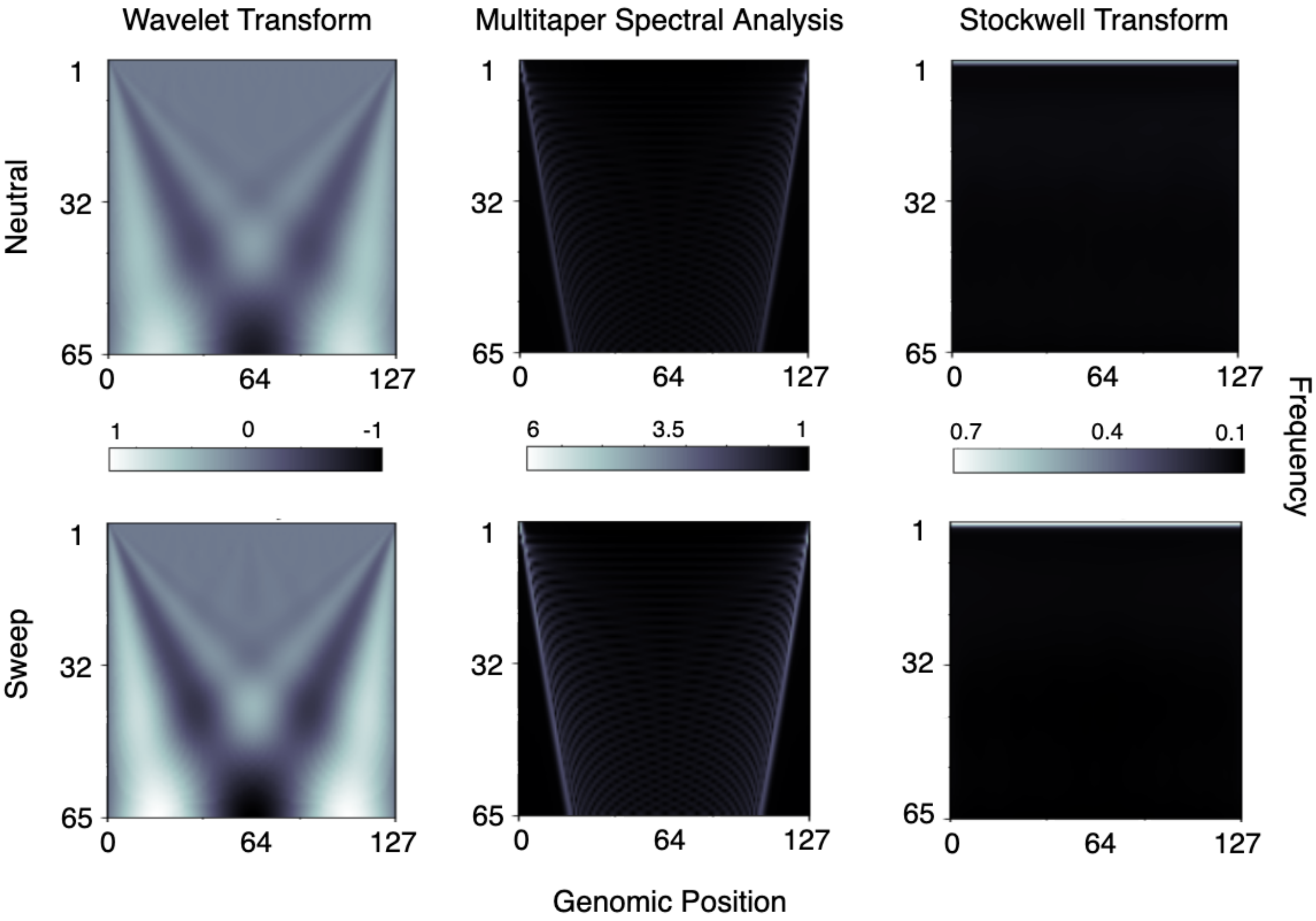


Figure S1: Mean time-frequency analysis input matrices for $n = 128$ windows of the mean pairwise sequence differences $\widehat{\pi}$ across the $N/2 = 9,000$ neutral and $N/2 = 9,000$ sweep replicates under the `Equilibrium_fixed` dataset containing an equilibrium constant-size demographic history and a sweep that completed $t = 0$ generations before sampling. Top row are neutral simulations and bottom row are sweep simulations. Time-frequency methods are depicted from left to right columns for the wavelet decomposition, multitaper analysis, and the S-transform, respectively.

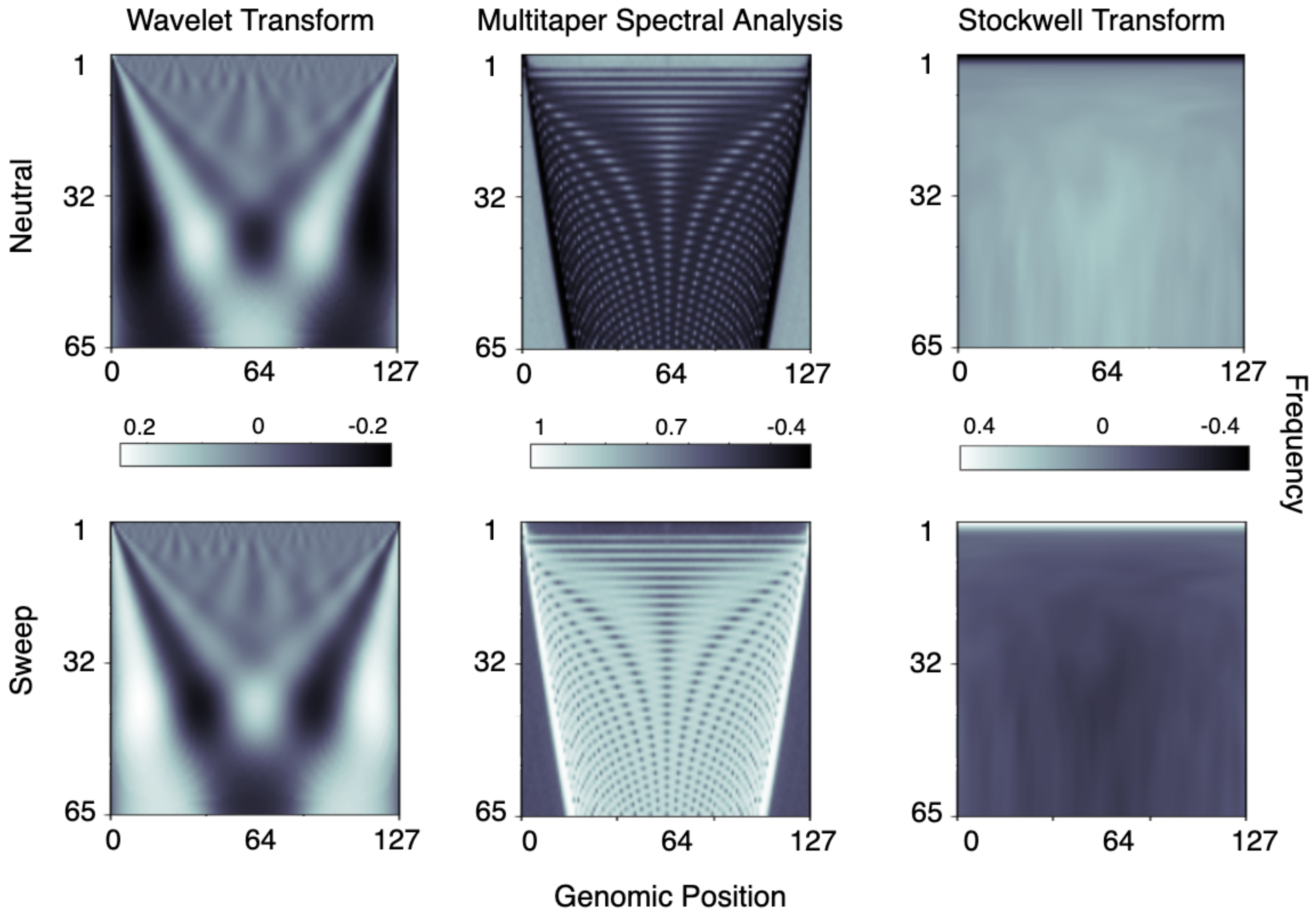


Figure S2: Mean time-frequency analysis input matrices for $n = 128$ windows of the mean pairwise sequence differences $\widehat{\pi}$ across the $N/2 = 9,000$ neutral and $N/2 = 9,000$ sweep replicates under the `Equilibrium_fixed` dataset containing an equilibrium constant-size demographic history and a sweep that completed $t = 0$ generations before sampling. Top row are neutral simulations and bottom row are sweep simulations. Time-frequency methods are depicted from left to right columns for the wavelet decomposition, multitaper analysis, and the S-transform, respectively. Elements of each matrix have been standardized to have a mean of zero and a standard deviation of one across all $N$ simulated replicates for a given time-frequency analysis method.

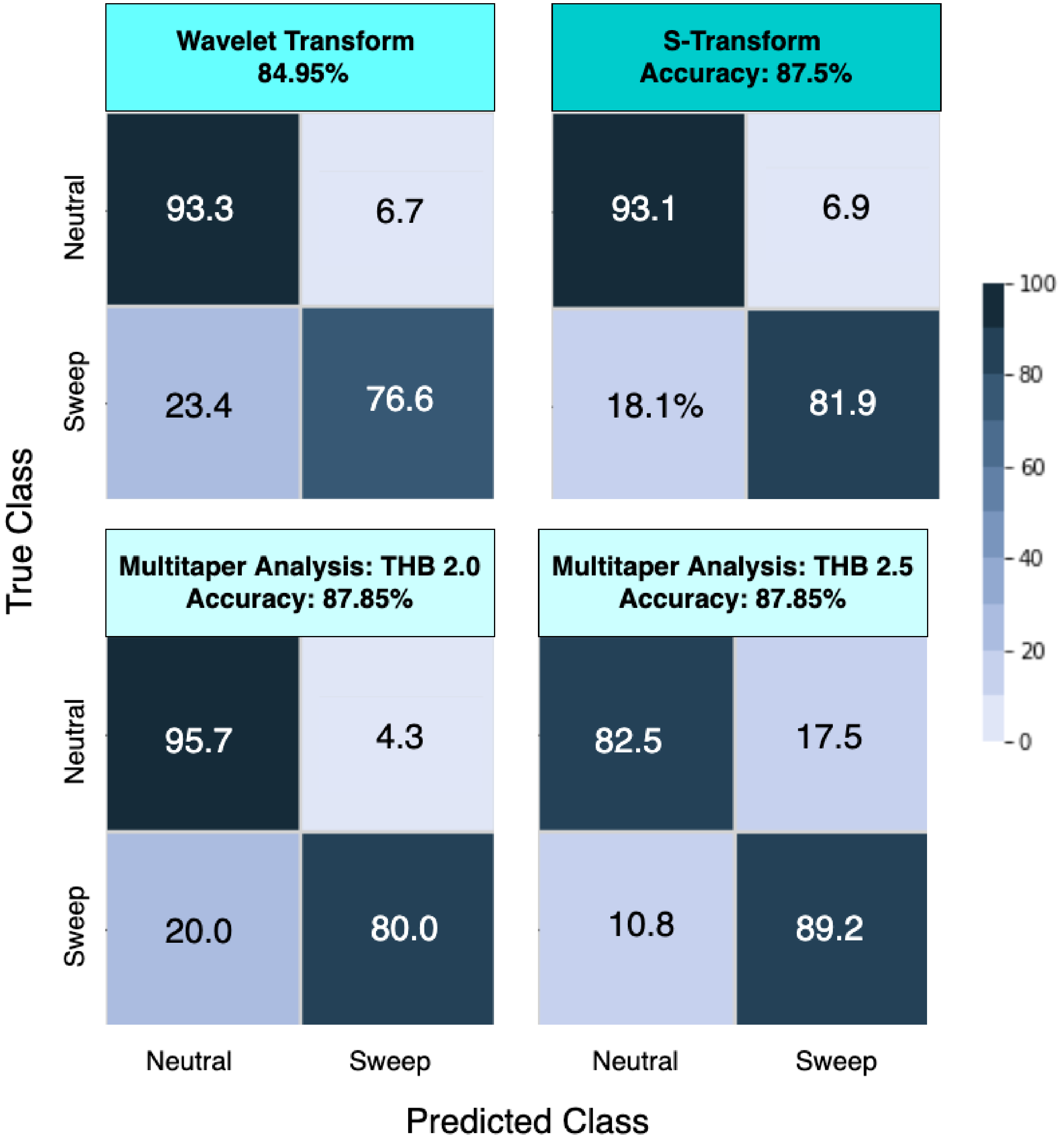


Figure S3: Classification rates and accuracy for a $c = 1$ channel convolutional neural network architecture (Figure 1) as depicted by confusion matrices for the `Equilibrium_fixed` dataset. Time-frequency analysis methods of wavelet decomposition, S-transform, and multitaper analysis with time-half bandwidth (THB) parameter of 2.0 and 2.5 were applied to an input signal of $n = 128$ windows of the mean pairwise sequence difference $\widehat{\pi}$.

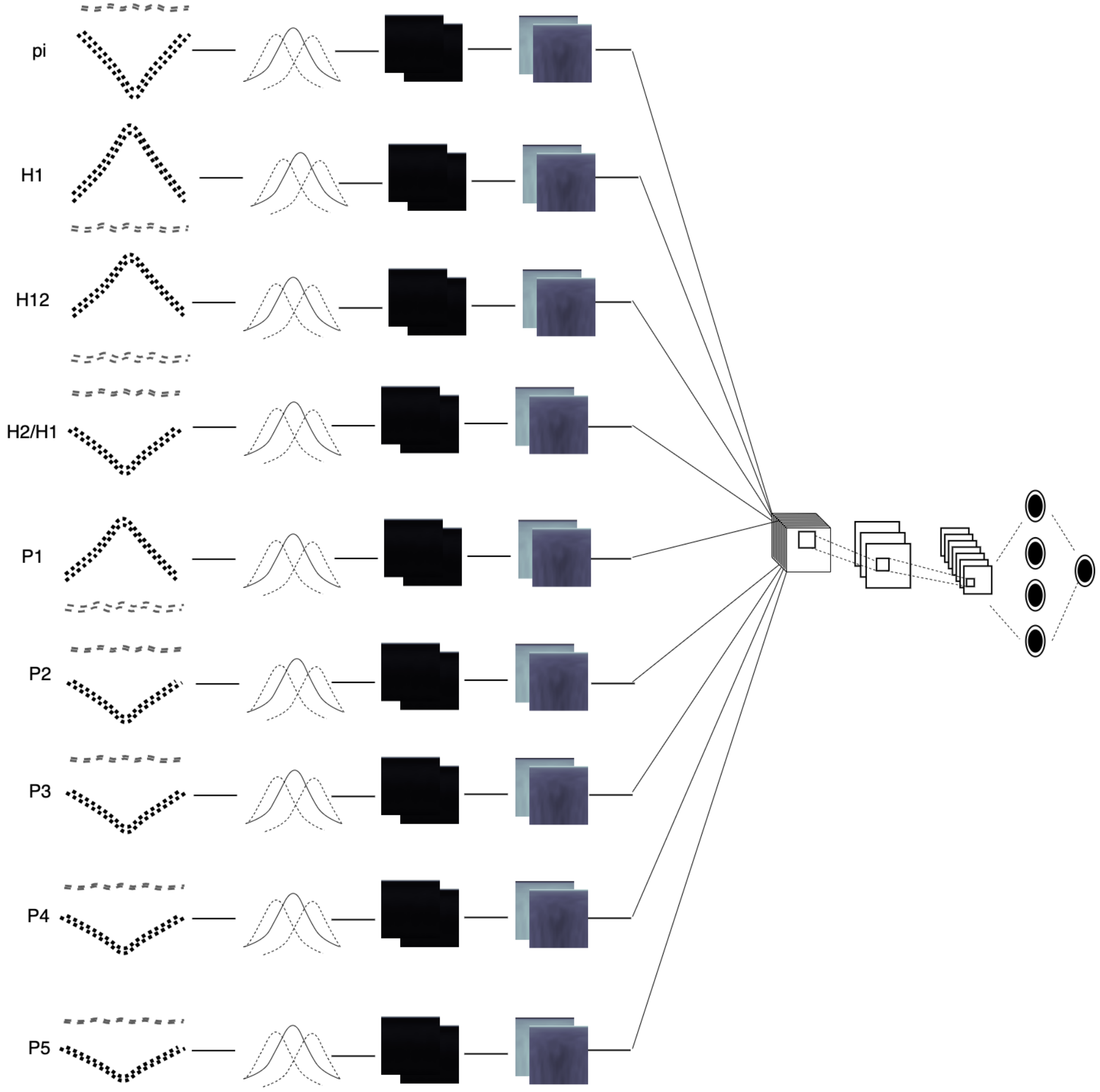


Figure S4: Depiction of a $c = 9$ channel convolutional neural network (CNN) architecture. Each summary statistic signal ($\widehat{\pi}$, $H_1$, $H_{12}$, $H_2/H_1$ and frequencies of the first five most common haplotypes respectively denoted by $P_1$ to $P_5$) of length $n = 128$ is used as input to a time-frequency analysis method (either wavelet decomposition, multitaper analysis, or S-transform) to decompose the signal into a matrix of dimensions $m \times n$, with $m = 65$, which is then standardized at each element based on the mean and standard deviation across all $N = 18,000$ training observations. These nine time-frequency analysis images are then stacked along a third dimension to create a tensor that is used as input to a CNN. The CNN has two convolution layers (three layers for the S-transform), followed by a dense layer with $n$ nodes containing both elastic-net and dropout regularization. The output layer of the CNN is a softmax that computes the probability of a sweep.

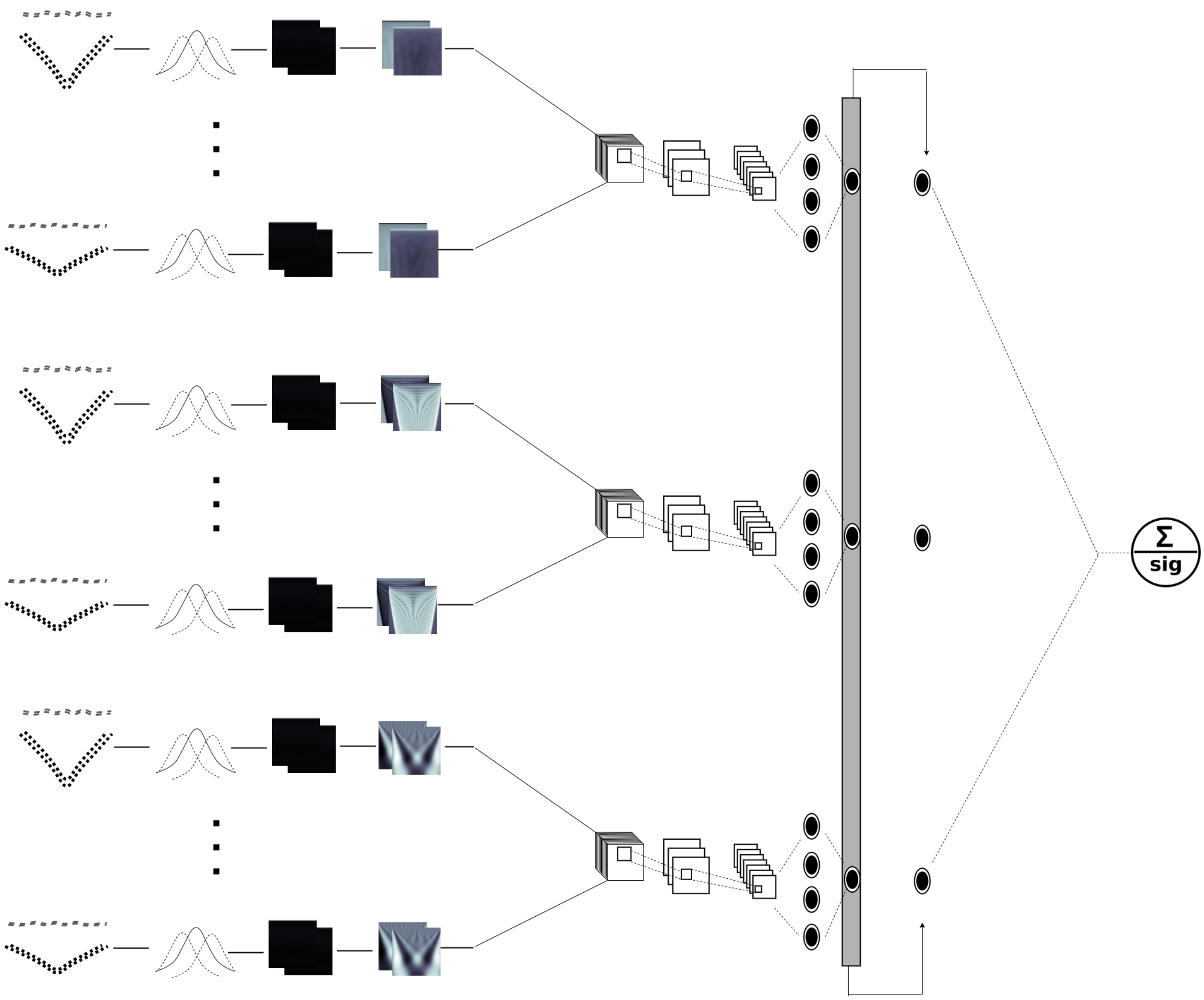


Figure S5: Depiction of the *SISSSCO*[*3CO*] model. Each summary statistic signal ($\widehat{\pi}$, $H_1$, $H_{12}$, $H_2/H_1$ and frequencies of the first five most common haplotypes respectively denoted by $P_1$ to $P_5$) of length $n = 128$ is used as input to each of the three time-frequency analysis method (wavelet decomposition, multitaper analysis, and S-transform) to decompose the signal into three matrices of dimension $m \times n$, with $m = 65$, which are then each standardized at each element based on the mean and standard deviation across all $N = 18,000$ training observations. For each time-frequency analysis method, the nine images (nine summary statistics) are stacked along a third dimension to create a tensor that is used as input to a CNN (as in Figure S4) and three independent $c = 9$ channel CNNs are trained for the three time-frequency analysis methods. The CNNs have two convolution layers (three layers for the S-transform), followed by a dense layer with $n$ nodes containing both elastic-net and dropout regularization. The output layer of the CNN is a softmax that computes the probability of a sweep. After training, the model parameters are fixed, and the output layers of the three CNNs are concatenated and these three nodes are used as input to a new output layer, which computes the probability of a sweep as a softmax.

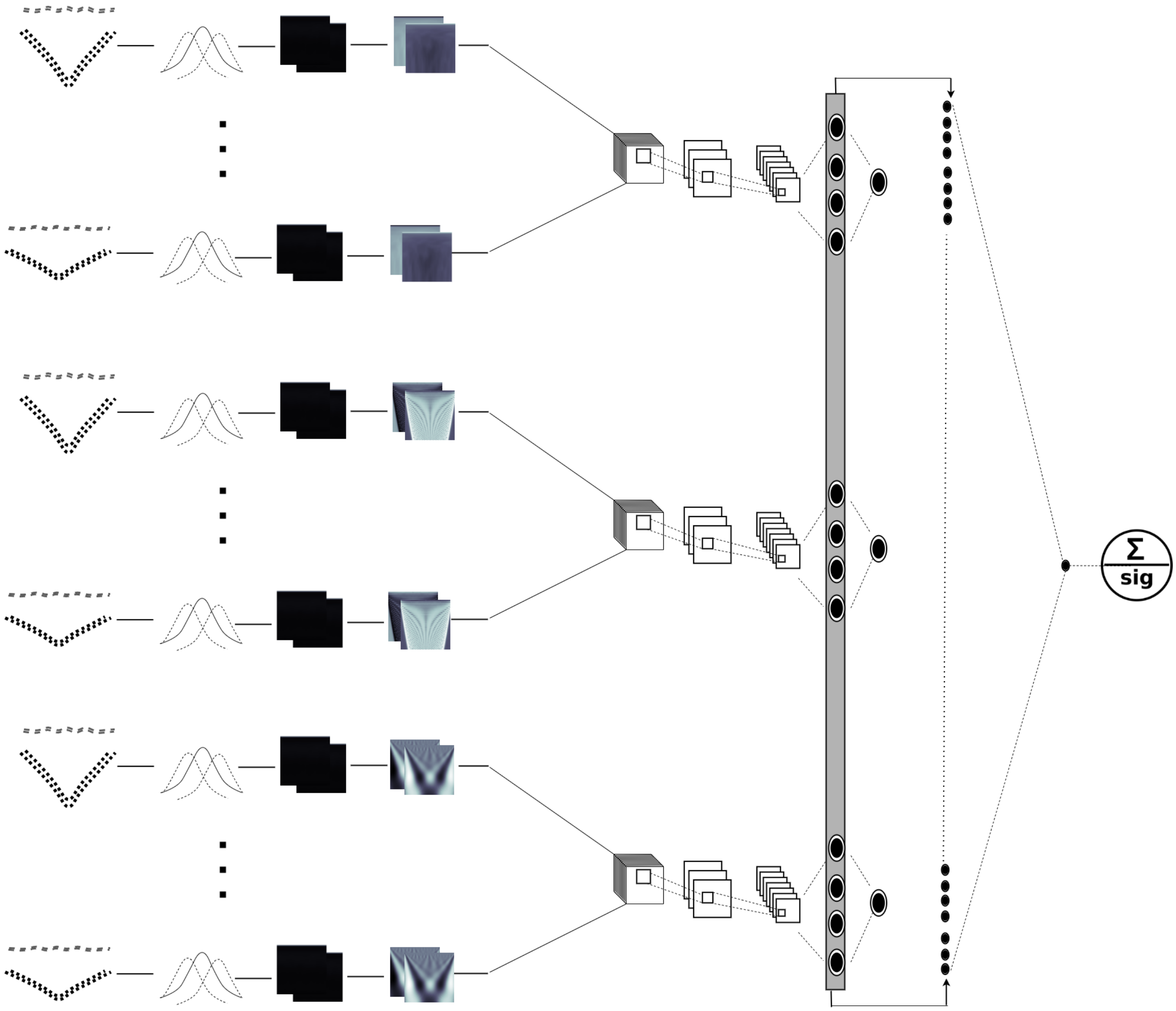


Figure S6: Depiction of the *SISSSCO*[*3CD*] model. Each summary statistic signal ($\widehat{\pi}$, $H_1$, $H_{12}$, $H_2/H_1$ and frequencies of the first five most common haplotypes respectively denoted by $P_1$ to $P_5$) of length $n = 128$ is used as input to each of the three time-frequency analysis method (wavelet decomposition, multitaper analysis, and S-transform) to decompose the signal into three matrices of dimension $m \times n$, with $m = 65$, which are then each standardized at each element based on the mean and standard deviation across all $N = 18,000$ training observations. For each time-frequency analysis method, the nine images (nine summary statistics) are stacked along a third dimension to create a tensor that is used as input to a CNN (as in Figure S4) and three independent $c = 9$ channel CNNs are trained for the three time-frequency analysis methods. The CNNs have two convolution layers (three layers for the S-transform), followed by a dense layer with $n$ nodes containing both elastic-net and dropout regularization. The output layer of the CNN is a softmax that computes the probability of a sweep. After training, the model parameters are fixed, and the dense layers of the three CNNs are concatenated and these $3n = 384$ nodes are used as input to a new output layer, which computes the probability of a sweep as a softmax.

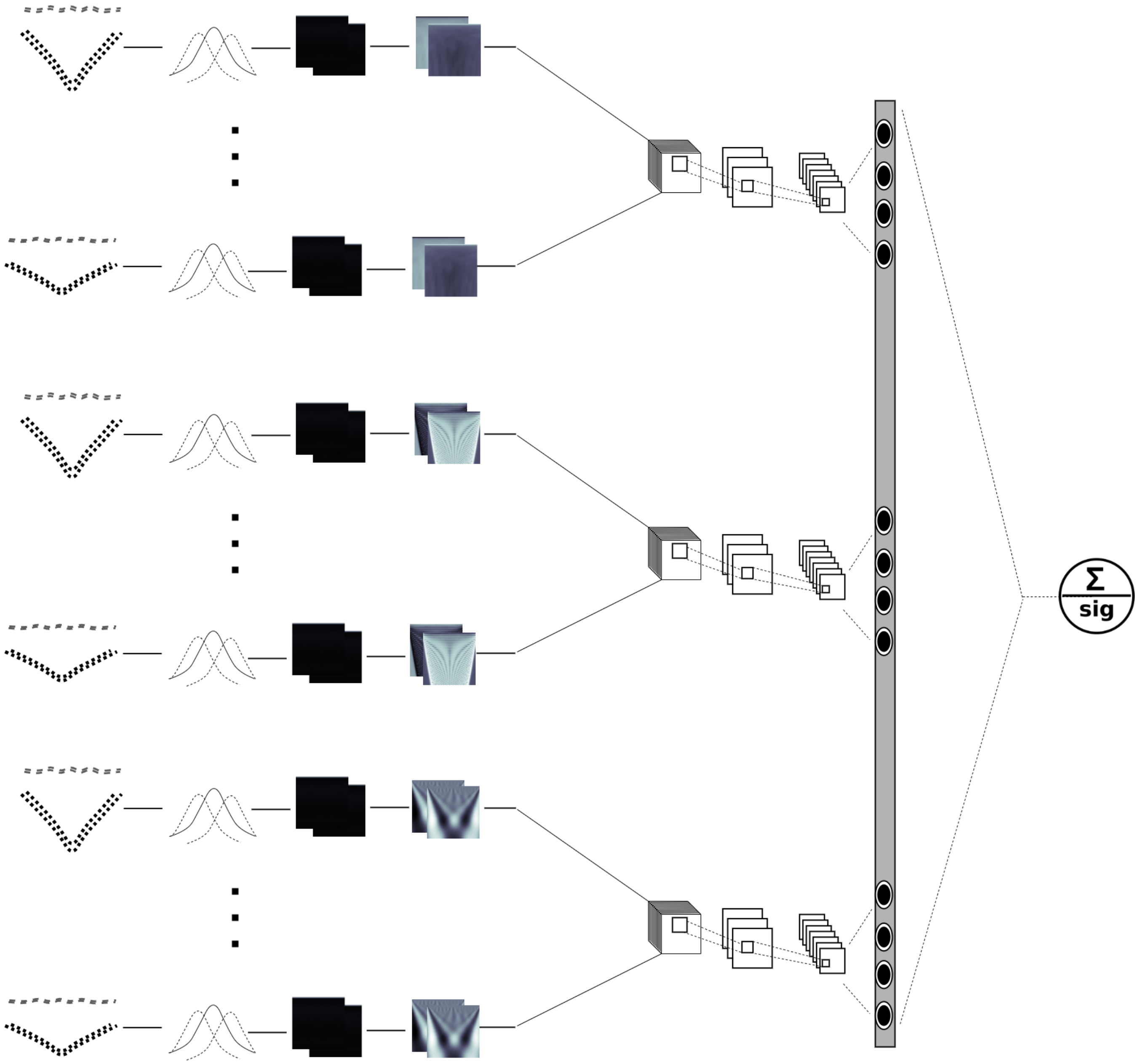


Figure S7: Depiction of the *SISSSCO*[*3MD*] model. Each summary statistic signal ($\widehat{\pi}$, $H_1$, $H_{12}$, $H_2/H_1$ and frequencies of the first five most common haplotypes respectively denoted by $P_1$ to $P_5$) of length $n = 128$ is used as input to each of the three time-frequency analysis method (wavelet decomposition, multitaper analysis, and S-transform) to decompose the signal into three matrices of dimension $m \times n$, with $m = 65$, which are then each standardized at each element based on the mean and standard deviation across all $N = 18,000$ training observations. For each time-frequency analysis method, the nine images (nine summary statistics) are stacked along a third dimension to create a tensor that is used as input to a CNN (as in Figure S4) and three independent $c = 9$ channel CNNs are trained for the three time-frequency analysis methods. The CNNs have two convolution layers (three layers for the S-transform), followed by a dense layer with $n$ nodes containing both elastic-net and dropout regularization. The $3n = 384$ nodes in the dense layers across the three CNNs are then fed into an output layer, which computes the probability of a sweep as a softmax. The parameters of the entire model of three CNNs are then estimated across the $N$ training observations.

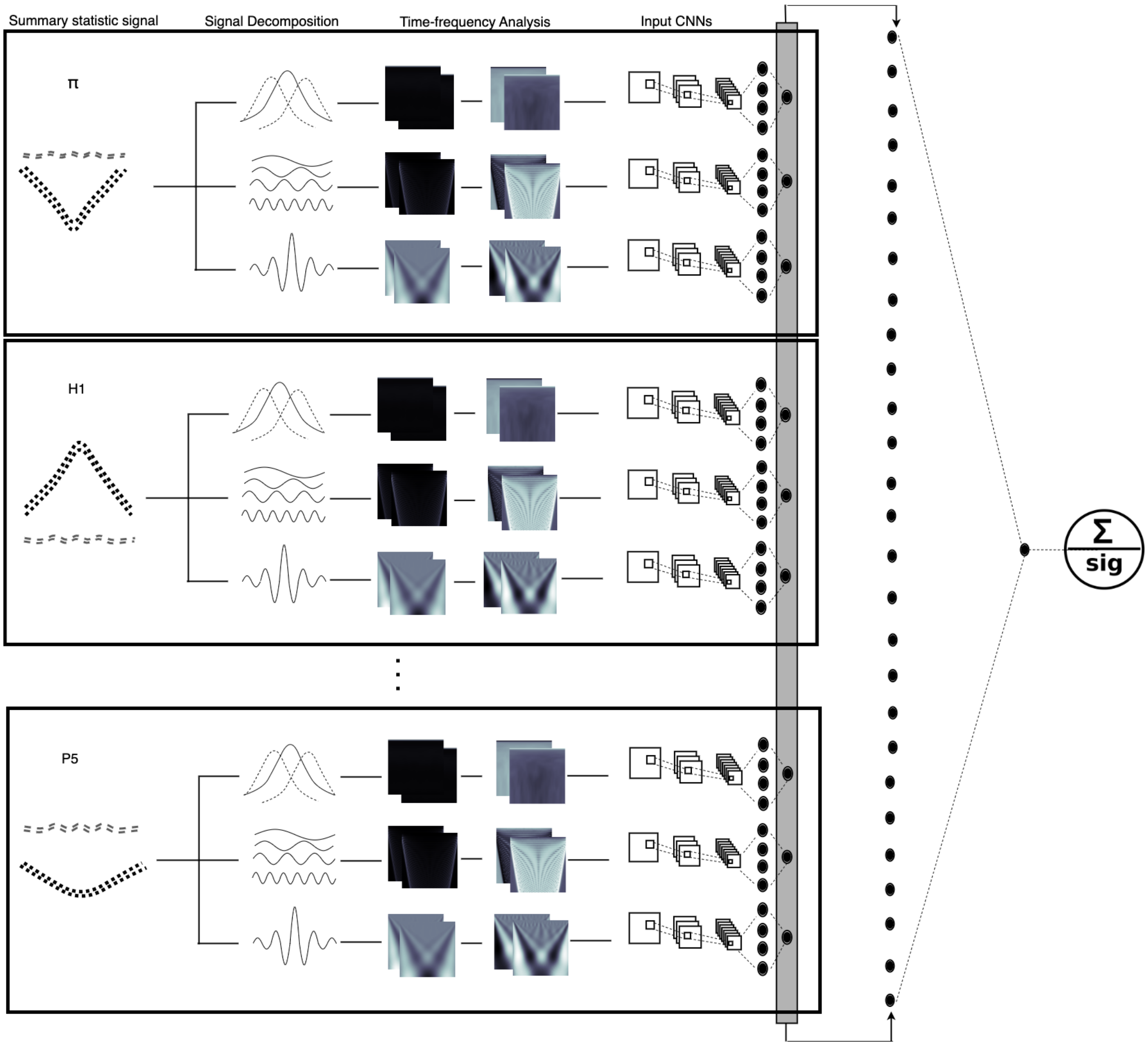


Figure S8: Depiction of the *SISSSCO*[*27CO*] model. Each summary statistic signal ($\widehat{\pi}$, $H_1$, $H_{12}$, $H_2/H_1$ and frequencies of the first five most common haplotypes respectively denoted by $P_1$ to $P_5$) of length $n = 128$ is used as input to each of the three time-frequency analysis method (wavelet decomposition, multitaper analysis, and S-transform) to decompose the signal into three matrices of dimension $m \times n$, with $m = 65$, which are then each standardized at each element based on the mean and standard deviation across all $N = 18,000$ training observations. These 27 images (nine statistics across three time-frequency analysis methods) each used as input to train 27 independent convolutional neural networks (CNNs). The CNNs have two convolution layers (three layers for the S-transform), followed by a dense layer with $n$ nodes containing both elastic-net and dropout regularization. The output layer of the CNN is a softmax that computes the probability of a sweep. After training, the model parameters are fixed, and the output layers of the 27 CNNs are concatenated and these 27 nodes are used as input to a new output layer, which computes the probability of a sweep as a softmax.

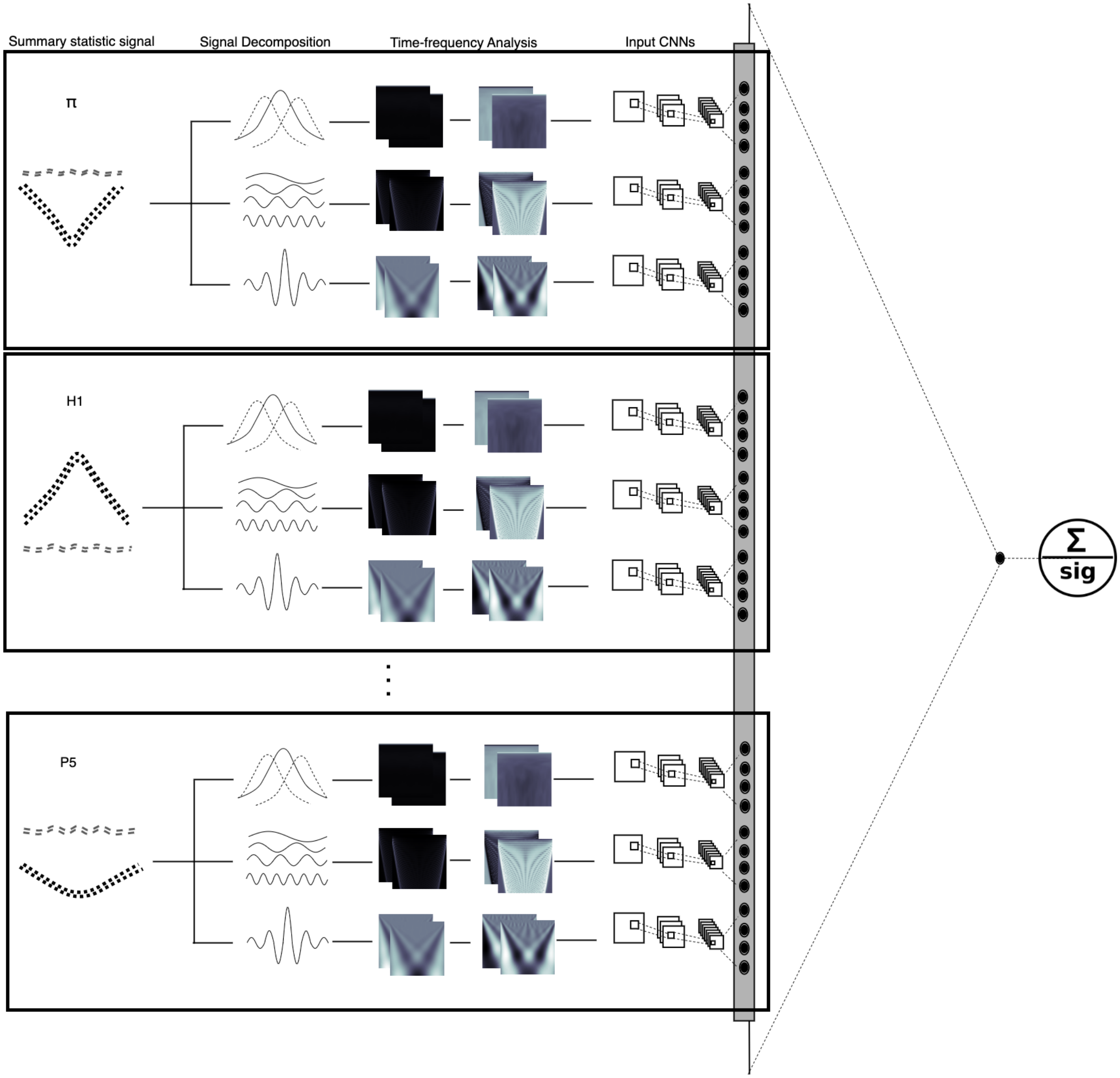


Figure S9: Depiction of the *SISSSCO[27MD]* model. Each summary statistic signal ($\hat{\pi}$, $H_1$, $H_{12}$, $H_2/H_1$ and frequencies of the first five most common haplotypes respectively denoted by $P_1$ to $P_5$) of length $n = 128$ is used as input to each of the three time-frequency analysis method (wavelet decomposition, multitaper analysis, and S-transform) to decompose the signal into three matrices of dimension $m \times n$, with $m = 65$, which are then each standardized at each element based on the mean and standard deviation across all $N = 18,000$ training observations. These 27 images (nine statistics across three time-frequency analysis methods) each used as input to 27 convolutional neural networks (CNNs). The CNNs have two convolution layers (three layers for the S-transform), followed by a dense layer with $n$ nodes containing both elastic-net and dropout regularization. The $27n = 3,456$ nodes in the dense layers across the 27 CNNs are then fed into an output layer, which computes the probability of a sweep as a softmax. The parameters of the entire model of 27 CNNs are then estimated across the $N$ training observations.

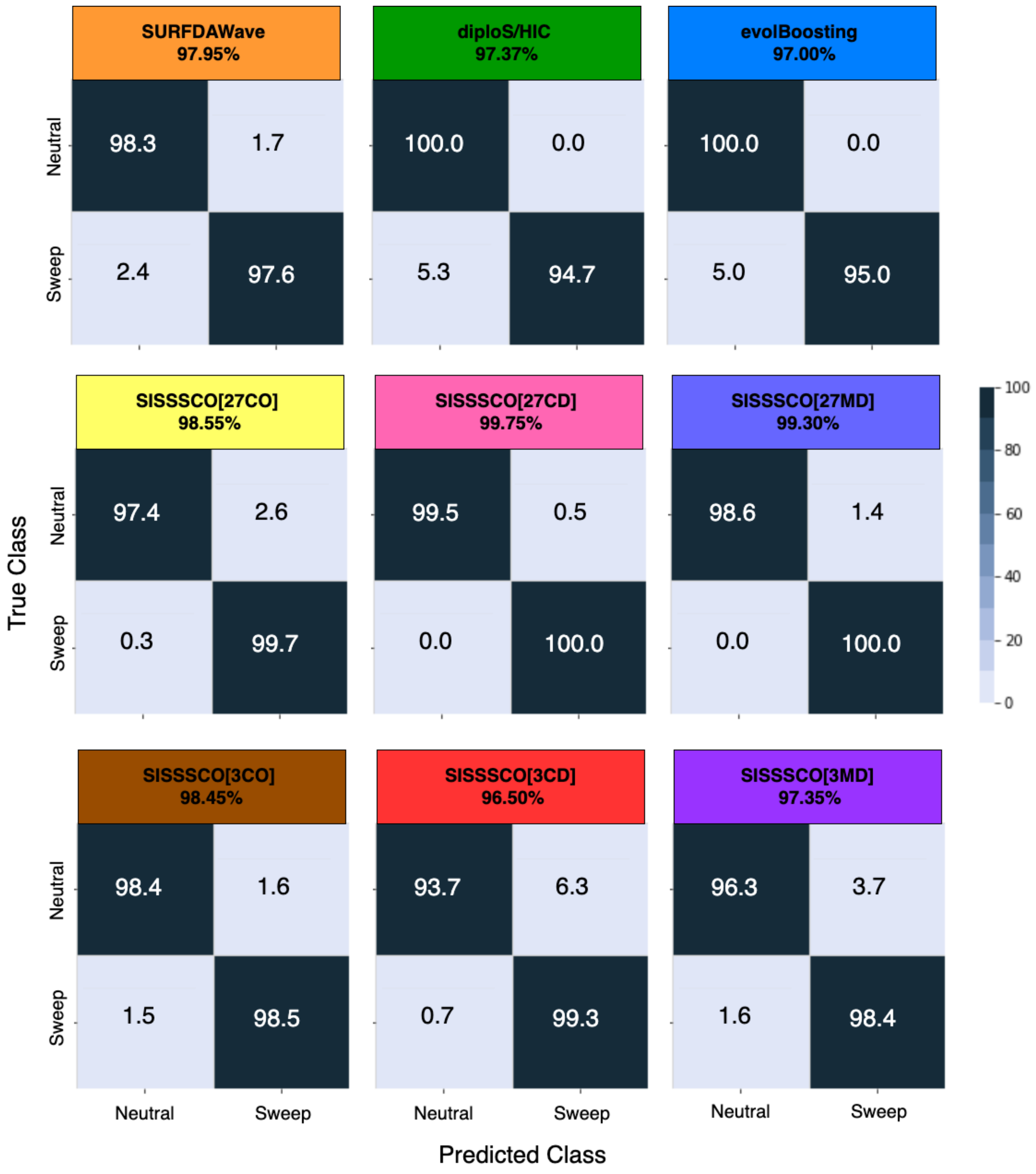


Figure S10: Classification rates and accuracy as depicted by confusion matrices to differentiate sweeps from neutrality on the `Equilibrium_fixed` dataset for the six *SISSSCO* architectures compared to *SURFDAWave*, diploS/HIC, and evolBoosting. The `Equilibrium_fixed` dataset is based on an equilibrium constant-size demographic history and a sweep that completed $t = 0$ generations before sampling.

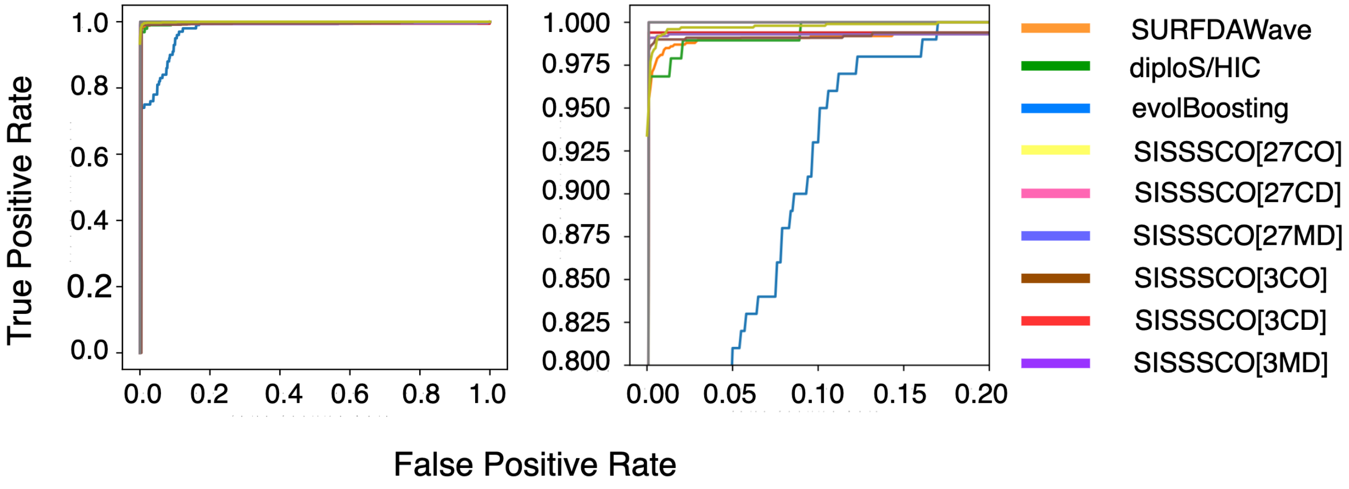


Figure S11: Power to detect sweeps as depicted by receiver operating characteristic curves on the `Equilibrium_fixed` dataset for the six *SISSSCO* architectures compared to *SURFDAWave*, diploS/HIC, and evolBoosting. The `Equilibrium_fixed` dataset is based on an equilibrium constant-size demographic history and a sweep that completed $t = 0$ generations before sampling. The right panel is a zoom in on the upper left-hand corners of the left panel.

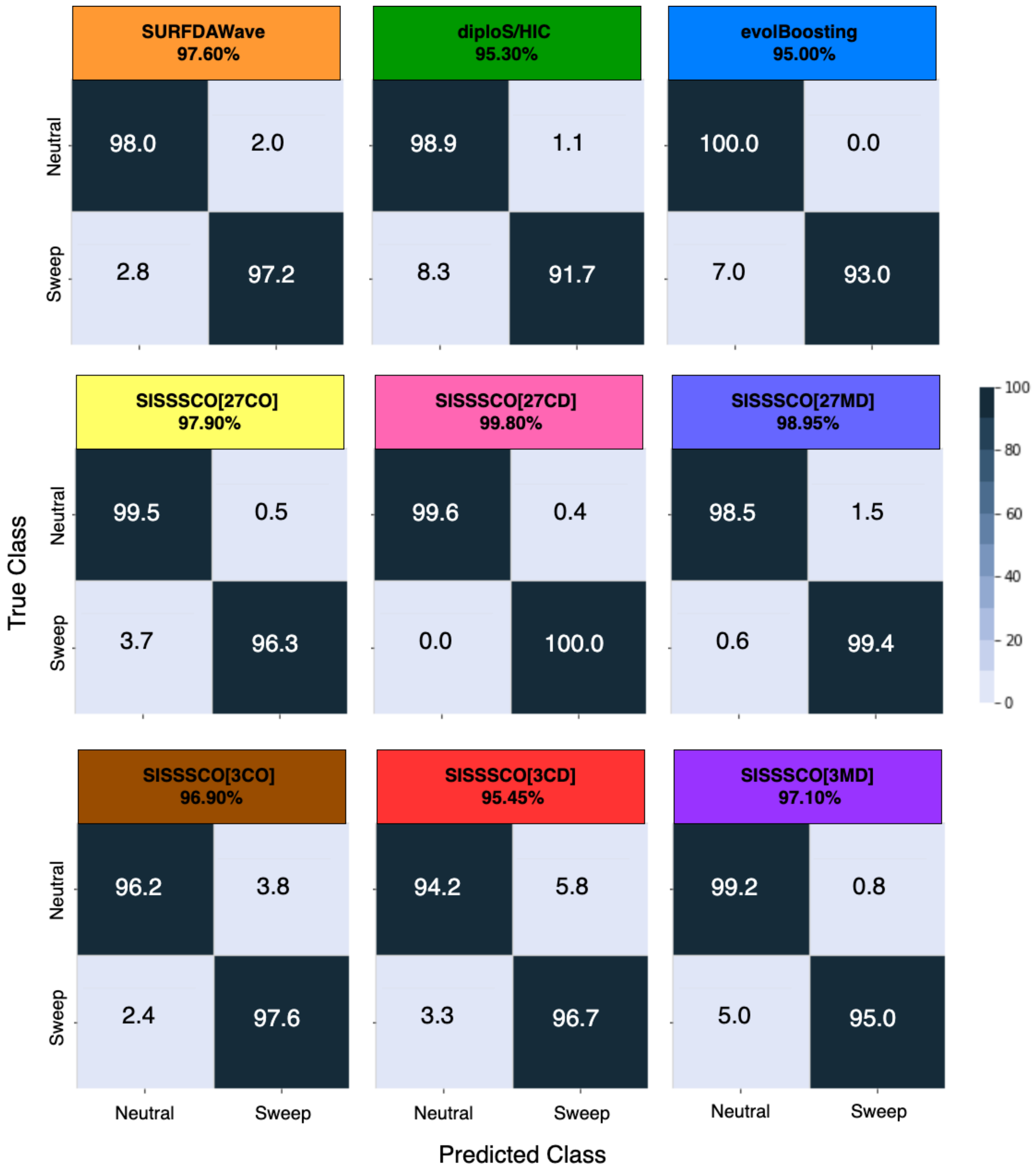


Figure S12: Classification rates and accuracy as depicted by confusion matrices to differentiate sweeps from neutrality on the `Equilibrium_variable` dataset for the six *SISSSCO* architectures compared to *SURFDAWave*, diploS/HIC, and evolBoosting. The `Equilibrium_variable` dataset is based on an equilibrium constant-size demographic history and a sweep that completed $t \in [0, 1200]$ generations before sampling.

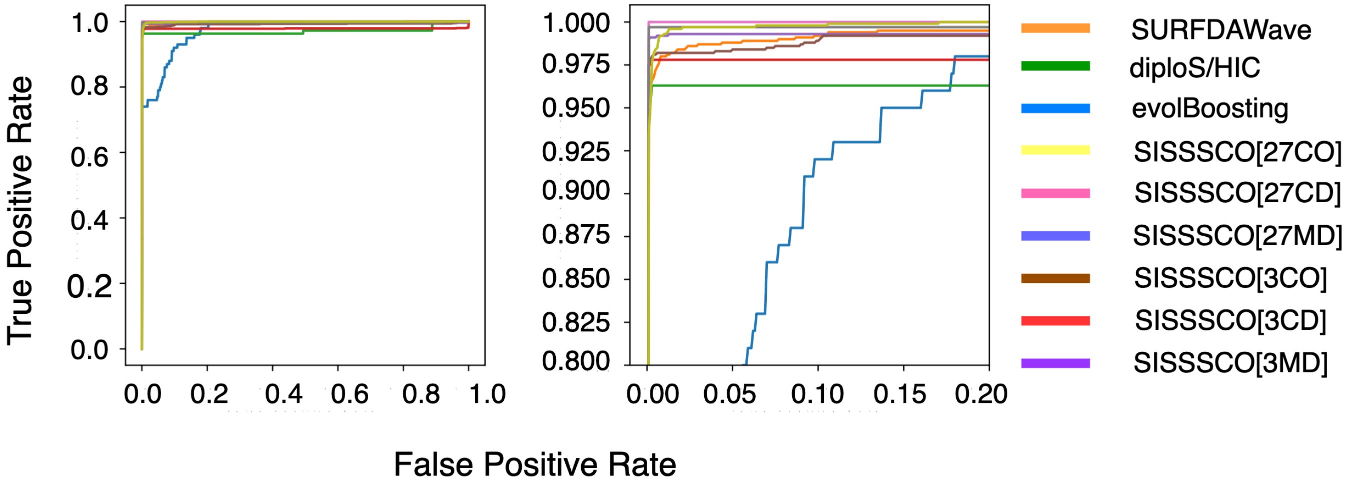


Figure S13: Power to detect sweeps as depicted by receiver operating characteristic curves on the `Equilibrium_variable` dataset for the six *SISSSCO* architectures compared to *SURFDAWave*, diploS/HIC, and evolBoosting. The `Equilibrium_variable` dataset is based on an equilibrium constant-size demographic history and a sweep that completed $t \in [0, 1200]$ generations before sampling. The right panel is a zoom in on the upper left-hand corners of the left panel.

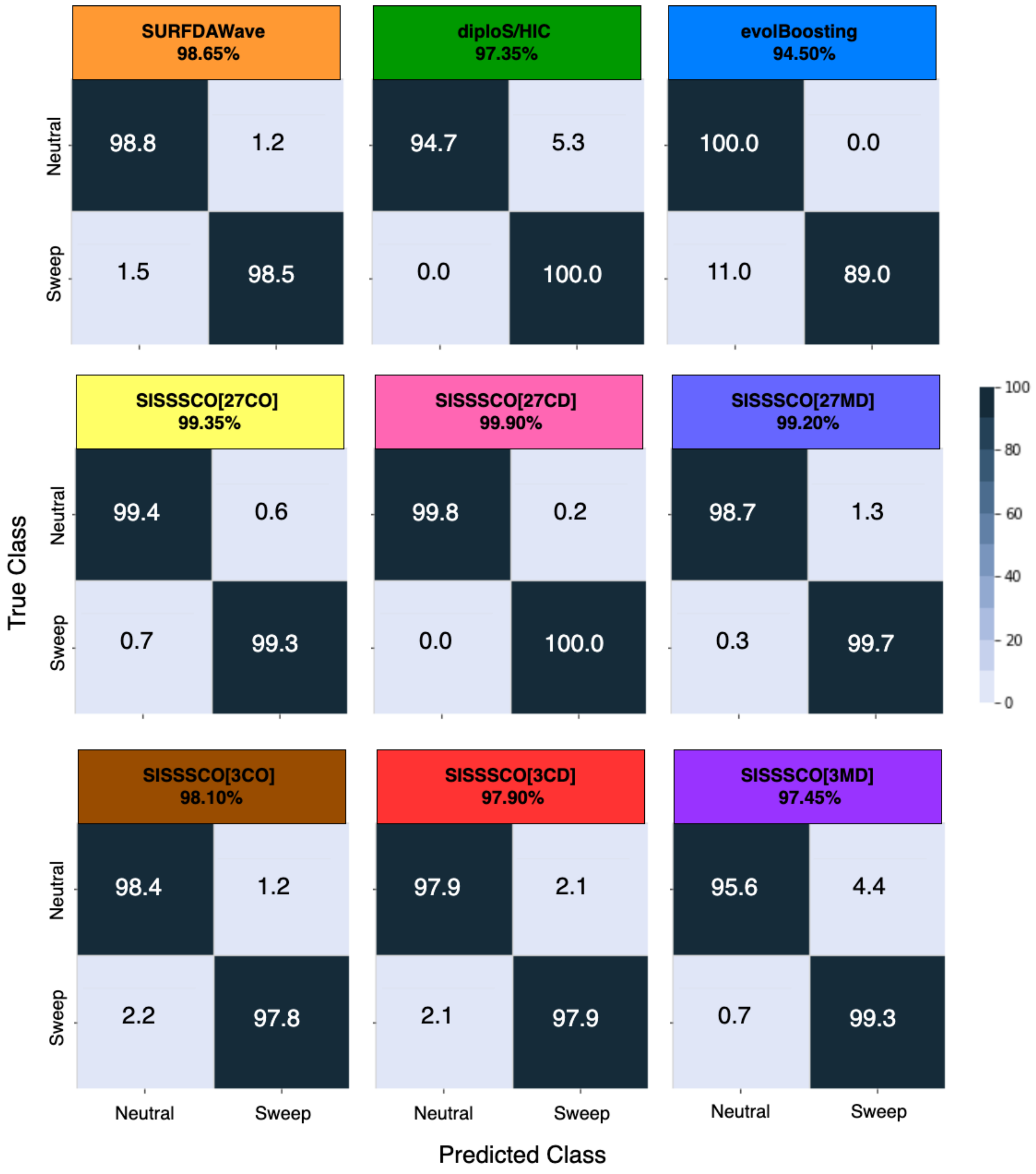


Figure S14: Classification rates and accuracy as depicted by confusion matrices to differentiate sweeps from neutrality on the `Nonequilibrium_fixed` dataset for the six *SISSSCO* architectures compared to *SURFDAWave*, diploS/HIC, and evolBoosting. The `Nonequilibrium_fixed` dataset is based on the nonequilibrium recent strong bottleneck demographic history of central European humans (CEU population in the 1000 Genomes Project) and a sweep that completed $t = 0$ generations before sampling.

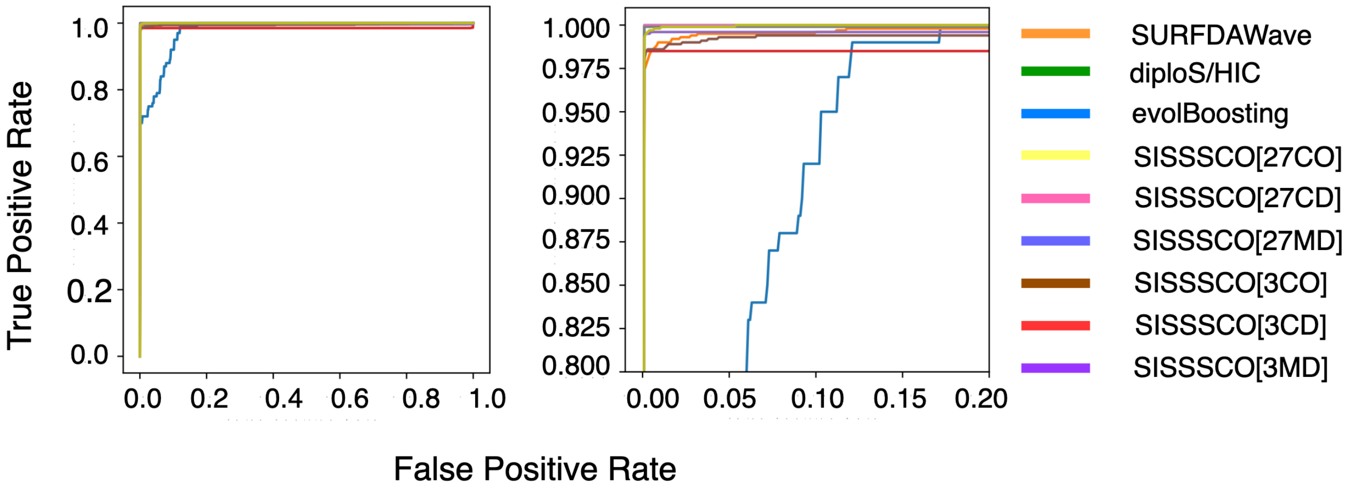


Figure S15: Power to detect sweeps as depicted by receiver operating characteristic curves on the `Nonequilibrium_fixed` dataset for the six *SISSSCO* architectures compared to *SURFDAWave*, diploS/HIC, and evolBoosting. The `Nonequilibrium_fixed` dataset is based on the nonequilibrium recent strong bottleneck demographic history of central European humans (CEU population in the 1000 Genomes Project) and a sweep that completed $t = 0$ generations before sampling. The right panel is a zoom in on the upper left-hand corners of the left panel.

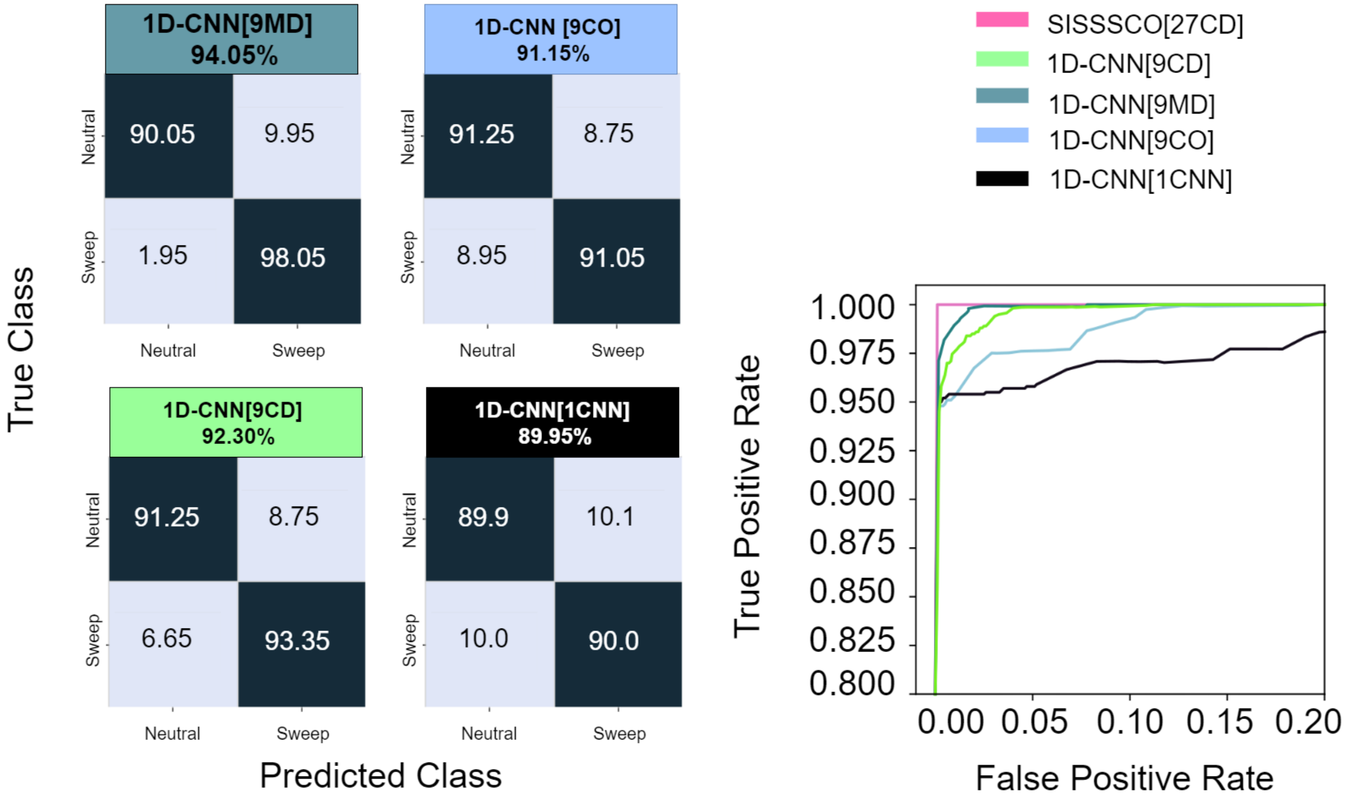


Figure S16: Right panel is the classification rates and accuracy as depicted by confusion matrices to differentiate sweeps from neutrality on the `Nonequilibrium_variable` dataset. The `Nonequilibrium_variable` dataset is based on the nonequilibrium recent strong bottleneck demographic history of central European humans (CEU population in the 1000 Genomes Project) and a sweep that completed $t \in [0, 1200]$ generations before sampling. Left panel is the power to detect sweeps as depicted by receiver operating characteristic curves on the `Nonequilibrium_variable` dataset for the *SISSSCO*[*27CD*] architecture compared to the *1D-CNN* architectures (*1D-CNN*[*1CNN*], *1D-CNN*[*9CO*], *1D-CNN*[*9CD*], and *1D-CNN*[*9MD*]). The right panel is a zoom in on the upper left-hand corner of the complete receiver operating characteristic curve.
